# Associations among Fish Length, Dam Passage History, and Survival to Adulthood in Two At-Risk Species of Pacific Salmon

**DOI:** 10.1101/572594

**Authors:** James R. Faulkner, Blane L. Bellerud, Daniel L. Widener, Richard W. Zabel

## Abstract

Threatened or endangered salmon and steelhead originating in the Snake River basin must pass through a series of eight major hydroelectric dams during their seaward migration. Understanding the effects of specific dam passage routes on lifetime survival for these stocks is essential for successful management. Juvenile fish may pass these dams via three primary routes: 1) spillways; 2) turbines; or 3) juvenile bypass systems, which divert fish away from turbines and route them downstream. Bypass systems may expose fish to trauma, increased stress, or disease. However, numerous studies have indicated that direct survival through bypass systems is comparable to and often higher than that through spillways. Some researchers have suggested that route of dam passage affects mortality in the estuary or ocean, but this is complicated by studies finding fish size affects route of passage. We tested whether passage through bypass systems was associated with probability of adult return after accounting for fish length and other covariates for two species of concern. We also investigated the association between fish length and probability of bypass at dams, and how this relationship could lead to spurious conclusions regarding effects of bypass systems on survival if length was ignored. We found that: 1) larger fish had lower bypass probabilities at 6 of 7 dams; 2) larger fish had higher probability of surviving to adulthood; 3) bypass history had little association with adult return after accounting for length; and 4) simulations indicated spurious effects of bypass on survival may arise when no true bypass effect exists, especially in models without length. Our results suggest that after fish leave the hydropower system, bypass passage history has little effect on mortality. Our findings underscore the importance of accounting for fish size in studies of dam passage or survival.

---

Four evolutionarily significant units from the Snake River Basin that are listed under U.S. Endangered Species Act - spring/summer Chinook salmon (*Oncorhynchus tshawytcha*), fall Chinook salmon, sockeye salmon (*O. nerka*), and steelhead trout (*O. mykiss*) - must pass a series of eight large hydroelectric dams, part of the Federal Columbia River Power System (FCRPS), on their migrations to the Pacific Ocean as smolts and upon their return as adults. A question central to the management of these populations is if the set of passage routes juvenile salmon take as they migrate through dams impacts their survival after they have completed their downstream migration.

Providing safe and effective downstream passage for juvenile salmon through the FCRPS has proven to be more problematic than adult upstream passage, which is achieved through the use of fish ladders. Juveniles have several possible routes to pass a dam (Figure 1). They can pass through spill, which is water passed directly through spill gates or through spillway weirs. Alternatively, juveniles can pass through the powerhouse where the hydroelectric turbines are located. However, at 7 out of 8 of the dams the majority of fish entering the powerhouse are diverted into a juvenile bypass system (JBS). Juvenile bypass systems are designed to divert fish in powerhouses away from turbine passage, using screens and a system of pipes that lead to a fish sampling and collection facility. From this facility, they can be directed into the dam tailrace, or (at three dams on the Snake River) loaded onto barges or trucks in a transportation program designed to avoid passage through downstream dams.

**Figure 1.**
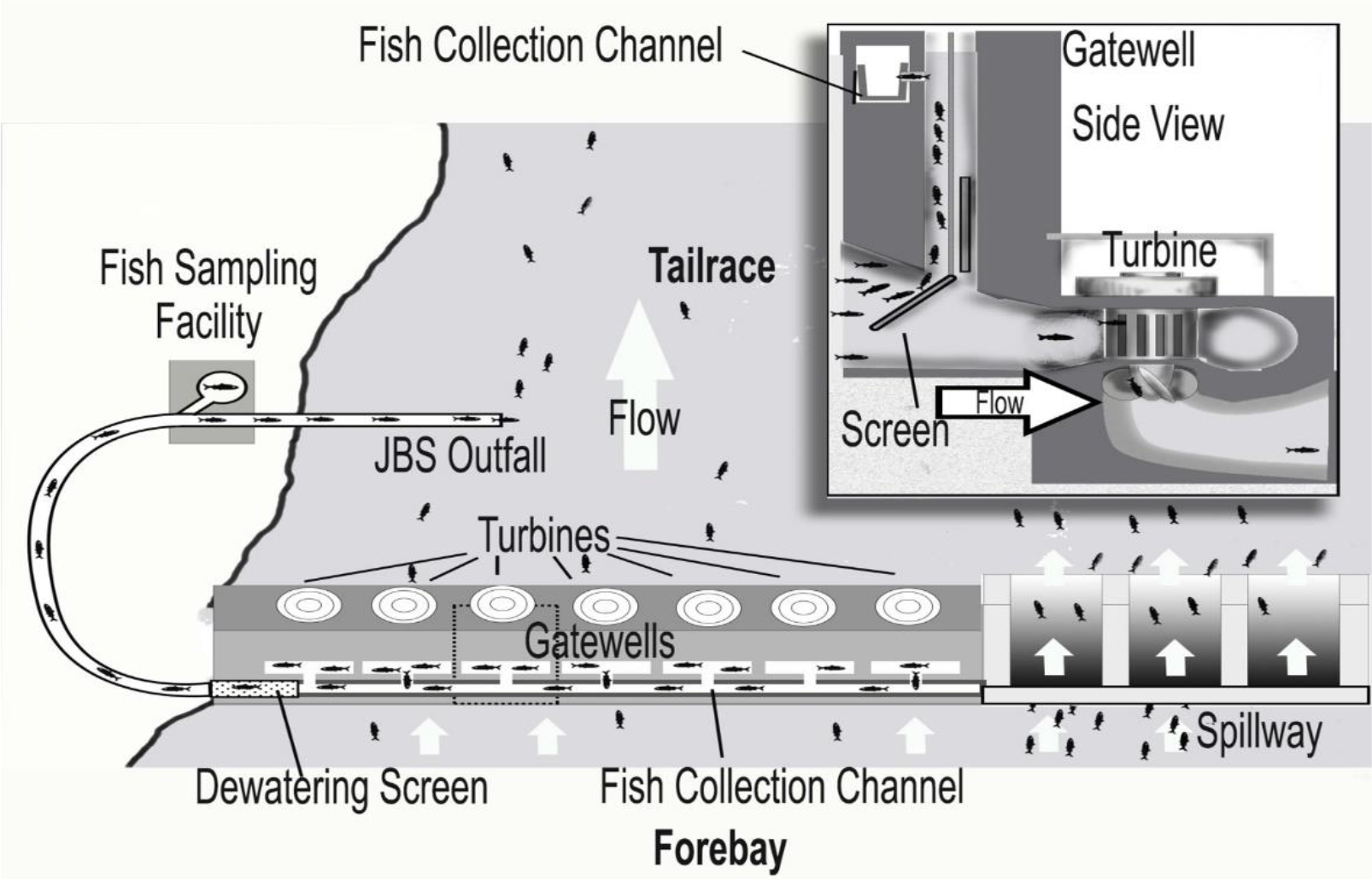
Overhead view of a hydroelectric dam with powerhouse and juvenile bypass system on the left and spillway on the right. Inset shows a turbine intake with bypass screen and entry into gatewell and collection channel for bypass system.

Numerous studies have been conducted with tagged fish to estimate survival through these various passage routes. Estimates of direct survival (to a short distance below the dam) from recent studies for yearling Chinook and steelhead ranged from 90-100% across eight dams, with a mean of 97% (Beeman et al. 2008; Axel et al. 2008; Ploskey et al. 2011; Ploskey et al. 2012; Skalski et al. 2013a; Skalski et al. 2013b; Weiland et al. 2013; Weiland et al. 2015). Differences between estimates of direct survival through spillway and JBS (spill - JBS) from these studies were relatively small, ranging from −5.4% to 5.0% with a mean of −0.8% for Chinook, and from −3.9 to 3.0% with a mean of −0.9% for steelhead, indicating slightly higher survival on average through JBS. Estimated probabilities of JBS passage from the previously mentioned studies ranged from 6.3 to 31.0% with a mean of 18.1% for yearling Chinook and from 5.9 to 41.9% with a mean of 22.1% for steelhead, while estimated probabilities of turbine passage at dams with a single powerhouse were much lower, ranging from 3.2 to 8.7% with a mean of 5.4% for yearling Chinook and from 1.8 to 5.8% with a mean of 3.2% for steelhead.

Although direct JBS passage survival is relatively high, there is concern that there may still be long-term negative impacts on survival for fish passing through JBSs that is not detected by direct survival studies. Sandford and Smith (2002) found that smolt-to-adult return rates (SAR) were frequently lower for fish bypassed through a JBS one or more times in comparison to those never bypassed. This and other studies at the time led to the concept of delayed or latent mortality, which refers to mortality that occurs in the ocean or estuary based on stress, injury, or diminished condition experienced during downstream migration through the hydropower system. This idea was proposed by Schaller et al. (1999) and Deriso et al. (2001) in terms of effect of dam passage in general. A review by Budy et al. (2002) suggested that cumulative stress of passing through turbine or bypass systems might result in increased mortality risk downstream. Petrosky and Schaller (2010) and Schaller et al. (2013) also attempted to incorporate the effects of specific route of passage in their analyses. They found that ocean survival in Chinook was negatively correlated with an index of the expected number of times a group of fish pass through turbine or JBS routes and with longer travel times through the FCRPS. Haeseker et al. (2012) used an index of spill proportion experienced by groups of fish as a surrogate for individual routes of passage and found it was positively correlated with ocean survival. Buchanan et al. (2011) found that multiple bypass events for individual fish were associated with lower SAR in hatchery yearling Chinook and steelhead from the Snake River basin, but timing and location of mortality could not be estimated.

Another possible explanation for reduced SARs of bypassed fish is that smaller fish and those in poorer condition tend to enter bypass systems with higher probability (Zabel et al. 2005; Hostetter et al. 2015), and smaller fish and fish in poorer condition also have lower SAR (Ward and Slaney 1988; Zabel and Williams 2002; Evans et al. 2014). This suggests that perceived effects of bypass on SAR could be at least in part due to correlation between bypass probability and fish size and condition and not due to bypass passage itself.

Even though numerous authors have demonstrated a difference in SAR between fish that experienced one or more bypasses compared to those never bypassed, success in demonstrating a cause and effect relationship has been limited. One obvious causal mechanism is that impaired function due to injury or stress caused by JBS passage could make smolts more susceptible to predation. Hostetter et al. (2012) found that steelhead with injuries or disease were more likely to be predated on by piscivorous birds in the Columbia River estuary. Although Budy et. al (2002) and other authors discuss the potential for injury passing through the pipes and over the screens of the bypass system, recent descaling and injury rates in bypass systems on the Snake and Columbia rivers are minimal (Ferguson et al. 2005), and much lower than those in the 1970s and early 1980s (Williams and Matthews 1995). Indeed, Sandford et al. (2012) found no significant difference in survival between bypassed and non-bypassed fish in seawater challenge experiments conducted on fish surviving passage through the FCRPS, suggesting that there was no bypass effect on subsequent survival.

Our main objective for this study was to investigate potential residual or lasting effects of bypass passage on post-hydropower system survival. We used an extensive dataset on juvenile Chinook and steelhead tagged at or upstream of Lower Granite Dam, the first dam encountered during downstream migration. We attempted to isolate lasting effects of bypass by only using fish known to have survived to downstream of the final dam in the hydropower system. However, we also recognized the need to account for fish size in our investigations, given established associations between fish size and probability of bypass and probability of adult return. In what follows, we first investigate the relationship between fish size at tagging and the probability of passing through a juvenile bypass system. We then investigate the association between fish length and SAR. We found that fish with more bypass events tend to have lower SAR, but evidence of a causal effect of bypass passage is greatly diminished or disappears when fish length is accounted for. Using simulated data, we also found that associations between bypass probability and fish length can lead to spurious estimates of negative effects of bypass passage on SAR when length is not accounted for.

## Methods

### Data

We used data on tagging and detection history for spring-summer Chinook and steelhead originating in the Snake River basin and implanted with passive integrated transponder (PIT) tags (Prentice et al 1990a) as juveniles. PIT tags are used extensively in salmon research because they are the only tagging medium that allows unique identification of individual fish from juvenile through adult life stages. We downloaded the PIT-tag data from the PTAGIS database (PSMFC 2017). Tagging data included locations and times of tagging and release, rearing type (hatchery or wild), and information about the researcher and study associated with each individual fish. We restricted the data to include only those fish with fork length (mm) recorded at time of tagging. Detection data included the location and time of detection of individual fish at any site with PIT-tag detection systems (Prentice et al 1990b), providing information from both juvenile and adult life stages.

Detection of PIT-tagged juveniles is possible at seven of the eight hydroelectric dams on the lower Snake and lower Columbia Rivers (Figure 2). The Dalles Dam (river kilometer (rkm) 308 from the ocean) is the only one of these dams without juvenile detection. Lower Granite Dam (LGD; rkm 695) is the furthest upstream dam and Bonneville Dam (BVD; rkm 234) is the furthest downstream in the FCRPS. The seven dams with tag detection have detectors installed in the JBSs, but BVD has additional detection in a sluiceway known as the corner collector (BCC). The BCC is located next to one of the two powerhouses and is designed to pass fish via water collected from the surface and directed through a gently sloping flume for several hundred feet to the tailrace. The final detection site for juveniles is in the Columbia River estuary in a detection array towed behind a pair of boats near rkm 50 (Ledgerwood et al. 2004). We will refer to this detection site as the estuary towed array (ETA). The main sites of detection of adults during our study were in fish ladders at BVD, McNary Dam (MCD; rkm 470), Ice Harbor Dam (IHD; rkm 538), and LGD.

Lower Granite Dam is the first dam encountered by fish migrating from upstream and large numbers of fish are tagged at the dam each year for research studies, with large proportion of those also measured at tagging. Although many fish tagged upstream of LGD are measured, tagging can occur from weeks to several months before those fish enter the hydropower system. Substantial growth can occur between tagging and entering the hydropower system. We therefore restricted our analyses to include fish tagged at LGD and those tagged at sites close enough upstream of LGD to reach LGD in less than 3 weeks on average. We further restricted the date of tagging to be between 18 March and 30 June, and the date of release to be between 1 April and 30 June. This restricted the set of fish to spring migrants and reduced the expected amount of growth before passing LGD. Juvenile bypass systems are turned off in the winter months and are typically turned back on around 1 April. Restricting release date to 1 April or later ensured all fish had an opportunity for detection. The combination of date and travel time restrictions resulted in the Snake River Trap (rkm 747), Clearwater River Trap (rkm 756), and Grande Ronde River Trap (rkm 795) being the only tagging and release sites upstream of LGD.

**Figure 2.**
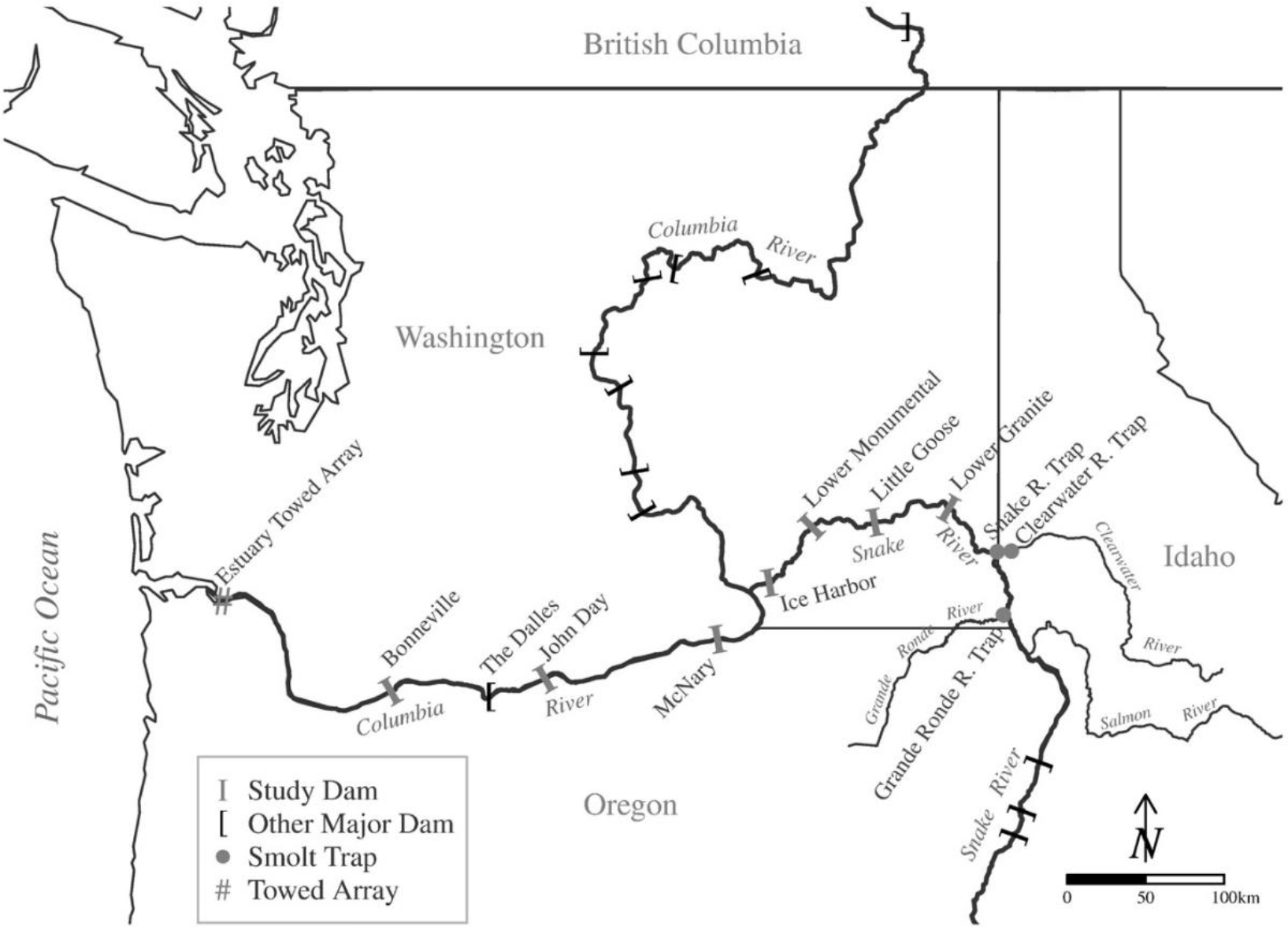
Study area showing locations of rivers, dams, the estuary towed array, and smolt traps used as tagging and release sites upstream of Lower Granite Dam. Study dams are those dams on the Snake River or lower Columbia River with PIT tag detection.

We performed two different main analyses: one investigating the association between length and probability of bypass, and the other investigating the associations of length and bypass history with probability of returning as an adult (SAR). Additionally, we investigated the association between length and detection in the BCC in comparison to the JBS at BVD. For all analyses, we used only fish detected either at BVD or ETA as juveniles, and therefore known to be alive while in the hydropower system. Any mortality experienced by these fish thus occurred after passage through all of the dams.

For comparisons between length and number of bypass events, we used fish detected as juveniles at BVD or ETA in the years 2000-2014. This set of years was chosen to be consistent with those used for adult return models (see below), but allowed a few additional years to increase sample sizes. Juvenile detections were not possible at IHD until 2005, so data for models there were restricted to 2005-2014. Juvenile detection was not possible in the BCC at BVD until 2006, so fish detected as juveniles at BVD prior to that passed through the bypass system at BVD. For specific comparisons between bypass and corner collector passage at BVD we used data from 2006-2014.

For SAR modeling, we used fish detected as juveniles at BVD or ETA in the years 2004- 2014. Equipment for PIT tag detection of adults at BVD was not consistent with modern configurations until 2004 and sample sizes in some years prior to that were low, so we excluded years prior to 2004. The cutoff of 2014 allowed nearly complete returns at the time of data acquisition in early 2017. Any fish detected in a fish ladder (adult fishway) at one or more dams in any year after its juvenile migration year was known alive at BVD and recorded as an adult return. Therefore, we defined SAR as survival from the site of last juvenile detection (BVD or ETA) to BVD as an adult. We excluded fish returning in less than one year (mini-jacks) from the SAR data because they may never reach the ocean and experience different conditions than fish returning after one or more years in the ocean.

For all data sets, we excluded fish tagged with acoustic or radio tags and those known to have been part of experiments involving multiple handling events or anesthetizations. We also excluded fish transported on barges, since these fish are known to have different survival and have fewer opportunities for detection than fish that remain in the river. Some summaries of the data are provided Appendix Tables A1 and A2.

### Modeling Bypass Probability

Our objective was to describe relationships between fish length and probability of passage through a JBS after accounting for other sources of variation. We used all fish with fork length measured at the time of tagging with restrictions described previously, where site of tagging was either LGD or sites upstream. We fit separate models for each dam, where the dams were: LGD, Little Goose Dam (rkm 635), Lower Monumental Dam (rkm 589), IHD, MCD, John Day Dam (JDD; rkm 347), and BVD. We used fish tagged upstream of LGD for modeling bypass probability at LGD. We used fish detected at ETA for modeling bypass probability at BVD. For all other dams, we used the combined set of fish tagged at or upstream of LGD.

We modeled bypass probability with binary logistic regression, where a fish was given a 1 for JBS passage (bypassed) or 0 for passage through another route (not bypassed). We used three classes of model, which were distinguished by their representation of time in the season. The first used a categorical variable (*rperiod*) that grouped release days into three periods: 1) 1-21 April, 2) 22 April-12 May, and 3) 13 May-30 June. These dates resulted in similar numbers of fish in each period. The second model class used standardized day of passage at LGD (<u>*pday*</u>) and the third used standardized day of detection (*dday*) at BVD or ETA, where days of passage or detection were measured continuously to account for time of detection or passage. For fish tagged upstream of LGD, *pday* was the day and time of last detection at LGD if a fish went through the JBS, and was the release day and time plus the median travel time to LGD for the particular release site and year of the fish if it was not detected at LGD. For fish tagged at LGD, *pday* was the day and time of release. For the fullest possible models of each type, the probability (*p*_*i*_) of being bypassed at a dam for individual fish *i* was assumed to be a logit-linear function of the day variables, release site (*rsite*), and standardized length at tagging (*length*) with random intercepts by year and random slopes for date variables by year:

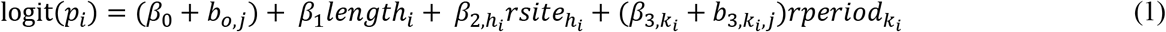

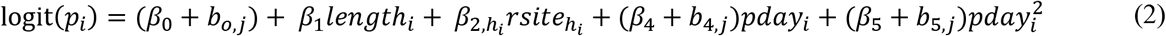

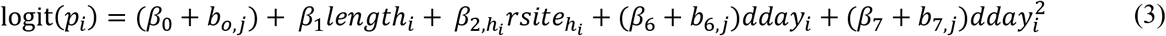

where *β*_0_ is the fixed intercept, *β*_1_, *β*_2,ℎ_, *β*_3,*k*_ and are the fixed effects of *length*, the *h*th level of *rsite*, and the *k*th level of *rperiod*, respectively. The fixed effects for the continuous *pday*, *pday*^2^, *dday*, and *dday*^2^ are represented by *β*_4_ through *β*_7_, respectively. The coefficients *b*_0,*j*_ and *b*_3,*j*_ through *b*_7,*j*_ are random effects for the migration year *j* in which fish *i* migrated. The sum of *β*_0_ + *b*_*o*,*j*_ amounts to a separate intercept for each year, and the sum of slope coefficients such as *β*_4_ + *b*_4,*j*_ allows the slopes to vary by year. The random effects were assumed to be independent and normally distributed with mean zero and a separate constant variance for each variable. The year and day effects and their combinations allowed us to account for variation in detectability at the dams due to river conditions and dam operations that vary with time but are difficult to measure and summarize for individual fish that are not detected. Models were constructed separately by species, rearing type, and dam.

We performed additional analyses for BVD that were a comparison between JBS and BCC passage only, which excluded spill and turbine routes. For these models we were able to use all fish that were detected in the JBS or BCC at BVD, whether or not they were detected at ETA. The purpose of these models was to test whether probability of passing through the JBS relative to the BCC was dependent on fish size. The response variable was 1 for fish entering the JBS and 0 for fish entering the BCC. We used the same set of explanatory variables as were used in the main analyses of bypass probability for BVD.

### Modeling Smolt-to-Adult Returns

Our objective was to test for associations between probability of returning as an adult (SAR) and bypass history after accounting for fish length and other factors that account for variation in SAR over time. We investigated three alternative variables to describe JBS passage (bypass) history: 1) a binary variable for detection in any JBS (yes/no), 2) a categorical variable with categories for 0, 1, 2, 3, or 4 or more bypass events, and 3) a continuous variable for number of bypass events. Data were from fish detected as juveniles at BVD or ETA and potential covariates included a categorical indicator variable for detection at ETA. Fish detected at both BVD and ETA were included only once in the data set, and the date at ETA was used for the detection date covariate for those fish. The time variables were a categorical variable for juvenile migration year and a continuous variable for day of year at either BVD or ETA, which were standardized separately for each location. Hatchery and wild fish were modeled together using an indicator variable for wild. We did not fit models separately by rearing type due to sample sizes, but we did fit models separately by species,

We also fit separate models for fish tagged at LGD and those tagged upstream of LGD. All fish tagged at LGD go through the JBS, but no non-bypassed fish are tagged or measured at LGD. This creates an unbalance in the data that could bias results if the fish tagged upstream of LGD were combined with those tagged at LGD. Also, fish tagged at LGD have lower SAR’s than those tagged upstream of LGD (unpublished data). For models with fish tagged upstream LGD we included a categorical variable for tagging site. Fish tagged upstream of LGD had a maximum of seven possible bypass events and those tagged at LGD had six, with the exception of 2004 when there were six and five possible, respectively, due to no detection at IHD. We combined data from the Clearwater River Trap with that from the Snake River Trap due to small sample sizes at the Clearwater River Trap and the close proximity of the two traps (10 km apart).

We assumed the binary variable for adult return followed a binomial distribution where the probability of return was a logit-linear function of the explanatory variables. For the fullest possible model, the logit of the probability of returning as an adult (*S*_*i*_) for individual fish *i* was:

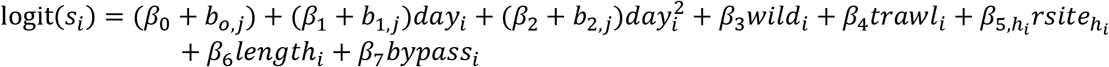

where *β*_0_ is the fixed intercept, and *β*_1_ through *β*_6_ are the fixed effects of standardized day at BVD or ETA, associated squared day, wild rear type, last detection at the trawl, *h*th level of release site, length at tagging, and bypass history, respectively. The bypass variable here is generic for one of the three possible bypass variables (binary, categorical, or continuous) and would have one level for either the binary or continuous versions and four levels for the categorical version. The random effects *b*_0,*j*_, *b*_1,*j*_, and *b*_2,*j*_ are associated with the migration year *j* for fish *i* for the intercept, *day*, and *day*^2^ variables, respectively. Similar to the models for bypass probability, the random effects allow separate values of the coefficients for the intercept, *day*, and *day*^2^ variables by year. The random effects were assumed to be independent and normally distributed with means zero and a separate constant variance for each variable.

We used the Akaike Information Criterion (AIC; Akaike 1973, Burnham and Anderson 2002) as a measure of the relative predictive ability of a set of competing models. For both the bypass and SAR analyses, we first constructed a set of models based on possible combinations of the fixed covariates and random effects that did not include the length or bypass variables (we will refer to these as covariate models). We selected the best model among those based on AIC, where lower AIC is better. We then added the variables of interest (length and/or bypass) to the best covariate models and recorded AIC and the *P*-values associated with the respective variables. Our objectives were to test whether the parameter estimates for the variables of interest were different from zero and to assess whether the variables of interest improved the predictive ability of the models. We interpreted the strength of evidence in favor of particular models or variables using a combination of the size of differences in AIC between models, associated Akaike model weights (Burnham and Anderson 2002), and the degree of *P*-values of individual effects. We used 95% confidence intervals to express uncertainty in parameter estimates. We used the `R` computing environment (R Core Team 2017) for all aspects of the analyses and specifically used the `R` package `lme4` (Bates et al. 2015) for fitting the generalized linear mixed models. The data and `R` code used in the analyses for this paper are available at: https://github.com/jrfaulkner/bypass-length-sar.

### Simulations

The association between fish length and probability of entering bypass systems makes it difficult to separate the individual effects of these variables on probability of returning as an adult. If length truly did have an association with SAR but number of bypass events did not, we wanted to know if number of bypass events would still come up as a significant predictor in SAR models due to its correlation with fish length. One way to address this question is by simulating data with both number of bypasses and SAR associated with length, but with no independent effect of number of bypass events on SAR. If number of bypass events then appeared as a significant predictor in models fit to such simulated data, where fitted models did not include length, that would suggest that an apparent bypass effect on SAR in real data could actually be explained by the association with fish length alone. Conversely, if the bypass effect was not significant in SAR models fit to the simulated data but was significant in models fit to real data, that would suggest that there was an effect of bypass passage which is separate from length, or there is an association between bypass and some other unmeasured variable which is also associated with SAR.

To address these questions, we used simulations to assess the chance of detecting a bypass effect on SAR if one did not actually exist. We did this by generating data from the best SAR models of observed data that contained length and other covariates but no bypass effects, then fit models with a term for number of bypasses to those simulated data and recorded the results. For each species and tagging location we generated 1,000 simulated data sets. We fixed the data for observed number of bypasses, length, and other measured covariates and only simulated adult return data for each fish. By fixing these covariates, the observed association between length and number of bypasses was preserved.

We simulated data by first drawing a set of model parameter values from a multivariate normal distribution where the mean was the vector of parameter estimates (both fixed and conditional random effects) from the best model with length and covariates (but no bypass effect) fit to the real data and the covariance matrix was the estimated joint covariance of the associated model parameters (see Appendix for details). Using the random draw of model parameters and the static covariate data, we then calculated predicted probabilities of adult return for each fish. We then simulated adult return data (0 or 1) for each fish by drawing from Bernoulli distributions given the set of predicted adult return probabilities. This process was replicated for a second set of simulations where the data generating model was the best covariate-only model within each species and tagging location. This second set of simulations was used for testing the effect of number of bypass events when there was no true association between SAR and bypass or SAR and length.

For each simulated data set, we fit three models. The null model (M0) contained just the fixed and random covariate terms. The second model (M1) added the term for number of bypass events and the third model (M2) added the length variable to M1. A fourth model (M3), which was M0 with length added, was also fit for comparison. For each model, we recorded the parameter estimate for number of bypasses and associated *P*-values and model AIC values. For M1 and M2, we recorded the proportion of simulations in which the parameter estimate for number of bypasses was negative and had a *P*-value < 0.05. For the model with number of bypasses and no length, we also recorded the proportion of simulations in which the parameter estimate for number of bypasses was negative and the AIC was lower than that of the null model. We also calculated the mean and 0.025 and 0.975 quantiles of the parameter estimates for number of bypasses across simulations.

If there were no association between number of bypasses and SAR, then we would expect the bypass variable to have a negative estimate approximately 50% of the time and we would expect it to be both negative and significant at the 0.05 level (two-sided test) approximately 2.5% of the time when models including the bypass variable were fit to SAR data simulated using a length effect but no bypass effect. These percentages are those expected by chance alone when there is no bypass effect on SAR. We would also expect that any apparent effect of bypass would be diminished when length was also in the fitted model.

## Results

### Bypass Probability

We found strong evidence at most dams that the probability of entering the JBS at a dam was negatively associated with fish length after accounting for the other variables for each species and rearing type (Table 1, Figure 2). The addition of the length variable to these models greatly reduced the model AIC (equivalently, resulted in small *P*-values for length) for each species and rearing type for all dams, with the exception of wild steelhead at LGD and neither species or rearing type at BVD when JBS passage was compared to all other routes (Table 1). However, when JBS passage was compared to only the BCC route at BVD, there was convincing evidence that probability of entering the JBS was negatively associated with length after accounting for other variables for all species and rearing type combinations except hatchery steelhead (Table 1, Figure 2). The strongest associations between bypass and length (based on magnitude of the parameter for length) occurred at MCD and JDD for both species and rear types, with the exception of wild Chinook at MCD.

**Table 1.** Estimated slope parameters and associated 95% confidence intervals, change in AIC (dAIC), and *P*-values for standardized length variables from models for bypass probability by species, rear type (hatchery (H) or wild (W)), and dam (see text for definitions). BVCC represents BVD when non-bypassed fish are from the corner collector (BCC) only. Parameter values represent the estimated change in log-odds of bypass at a dam associated with an increase of 1 standard deviation in standardized length, after accounting for other variables in the models. dAIC here is the change in AIC associated with adding the length variable to the best model without length at each dam. The sample size (*n*) is also shown for each data set.

| Species | Type | Dam | $n$ | Estimate | 95% CI | dAIC | $P$-value |
| --- | --- | --- | --- | --- | --- | --- | --- |
| Chinook | H | LGD | 4,062 | -0.121 | (-0.224, -0.018) | -3.3 | 0.0213 |
|  |  | LGSD | 62,176 | -0.205 | (-0.234, -0.175) | -182.8 | <0.0001 |
|  |  | LMD | 62,176 | -0.165 | (-0.199, -0.131) | -90.1 | <0.0001 |
|  |  | IHD | 61,026 | -0.216 | (-0.261, -0.170) | -85.7 | <0.0001 |
|  |  | MCD | 62,176 | -0.218 | (-0.245, -0.191) | -250.0 | <0.0001 |
|  |  | JDD | 62,176 | -0.360 | (-0.398, -0.322) | -352.4 | <0.0001 |
|  |  | BVD | 10,648 | 0.015 | (-0.090, 0.121) | 1.9 | 0.7774 |
|  |  | BVCC | 50,143 | -0.230 | (-0.259, -0.200) | -237.9 | <0.0001 |
|  | W | LGD | 4,169 | -0.168 | (-0.290, -0.045) | -5.2 | 0.0075 |
|  |  | LGSD | 26,217 | -0.216 | (-0.277, -0.155) | -46.7 | <0.0001 |
|  |  | LMD | 26,217 | -0.207 | (-0.274, -0.139) | -34.7 | <0.0001 |
|  |  | IHD | 15,578 | -0.198 | (-0.303, -0.092) | -11.6 | 0.0002 |
|  |  | MCD | 26,217 | -0.104 | (-0.158, -0.051) | -12.9 | 0.0001 |
|  |  | JDD | 26,217 | -0.249 | (-0.313, -0.185) | -56.0 | <0.0001 |
|  |  | BVD | 4,081 | 0.018 | (-0.172, 0.209) | 2.0 | 0.8491 |
|  |  | BVCC | 12,827 | -0.202 | (-0.275, -0.129) | -27.9 | <0.0001 |
| Steelhead | H | LGD | 8,290 | -0.125 | (-0.198, -0.051) | -9.1 | 0.0009 |
|  |  | LGSD | 31,608 | -0.156 | (-0.188, -0.125) | -90.9 | <0.0001 |
|  |  | LMD | 31,608 | -0.176 | (-0.213, -0.140) | -86.9 | <0.0001 |
|  |  | IHD | 24,433 | -0.192 | (-0.248, -0.136) | -42.4 | <0.0001 |
|  |  | MCD | 31,608 | -0.263 | (-0.305, -0.222) | -154.7 | <0.0001 |
|  |  | JDD | 31,608 | -0.292 | (-0.335, -0.248) | -169.6 | <0.0001 |
|  |  | BVD | 4,536 | -0.047 | (-0.217, 0.123) | 1.7 | 0.5890 |
|  |  | BVCC | 21,105 | -0.031 | (-0.075, 0.013) | 0.1 | 0.1725 |
|  | W | LGD | 2,746 | -0.093 | (-0.219, 0.033) | -0.1 | 0.1487 |
|  |  | LGSD | 20,801 | -0.192 | (-0.235, -0.150) | -77.2 | <0.0001 |
|  |  | LMD | 20,801 | -0.187 | (-0.234, -0.140) | -60.5 | <0.0001 |
|  |  | IHD | 12,642 | -0.203 | (-0.291, -0.115) | -19.1 | <0.0001 |
|  |  | MCD | 20,801 | -0.282 | (-0.333, -0.231) | -121.2 | <0.0001 |
|  |  | JDD | 20,801 | -0.316 | (-0.372, -0.259) | -126.7 | <0.0001 |
|  |  | BVD | 3,901 | -0.066 | (-0.243, 0.110) | 1.5 | 0.4604 |
|  |  | BVCC | 10,115 | -0.238 | (-0.307, -0.169) | -45.7 | <0.0001 |

For each species and rearing type, the best model for bypass probability that did not include length varied by dam, but all included random year effects for the intercept and most had random effects associated with the variables representing day in season (see Appendix Table A3).

### Smolt-to-Adult Returns

For Chinook tagged upstream of LGD, the best SAR model without length or bypass effects included fixed effects for rear type, release site, site of last detection, and day of year, and random year effects for the intercept (Appendix Table A2). Adding the length variable resulted in a decrease in AIC of 4.6 and *P* = 0.005, which provides moderate to strong evidence of an association between length and SAR (Table 3). When length was not in the model, adding the binary bypass variable increased AIC, while adding the categorical bypass variable decreased AIC by 1.5 with associated *P* = 0.039, and adding the continuous number of bypasses decreased AIC by 2.5 with *P* = 0.036. This suggests moderate evidence of an association between SAR and number of bypass events when length was not in the model. When length was in the model, adding the binary bypass variable resulted in an increase in AIC, but adding categorical bypass variable reduced AIC by 1.1 with *P* = 0.060 and adding the continuous variable for number of bypasses reduced AIC by 1.3 with *P* = 0.074. This suggests weak to moderate evidence of an association with number of bypasses after accounting for length. For the model with length and continuous number of bypasses, the odds of adult return increased by an estimated multiplicative factor of 1.35 for every increase of 1 standard deviation in length (16.1 mm; 95% CI: 1.07, 1.70), and the odds of return decreased by a multiplicative factor of 0.85 for each additional bypass event (95% CI: 0.71, 1.02; Figure 3).

**Table 2.** Results for SAR models by species and tag site, where tag sites are Lower Granite Dam (LGD) or sites upstream of LGD (ULGD). Also shown are number of individuals (smolts) and number of adult returns for each data set used for model fitting. Each row gives the terms in the model and the number of parameters (*np*), difference in AIC (ΔAIC) compared to the model with lowest AIC, the model Akaike weight (*w*), and *P*-values associated with respective length or bypass variables in the model. Each model had a set of covariates (*covs*) described in the text which were common to all models within a particular species and tag site. The bypass variables are binary bypass indicator (*byp*), categorical number of bypasses (*byp.cat*), and number of bypasses (*n.byp*).

| Species | Tag Site | Smolts | Returns | Model | $np$ | $\Delta AIC$ | $w$ | $P$-value | |
| --- | --- | --- | --- | --- | --- | --- | --- | --- | --- |
|  |  |  |  |  |  |  |  | length | bypass |
| Chinook | ULGD | 6,348 | 100 | $covs$ | 6 | 5.9 | 0.02 | - | - |
| | | | | $covs + len$ | 7 | 1.2 | 0.18 | 0.005 | - |
| | | | | $covs + byp$ | 7 | 7.1 | 0.01 | - | 0.349 |
| | | | | $covs + byp.cat$ | 10 | 3.9 | 0.05 | - | 0.039 |
| | | | | $covs + n.byp$ | 7 | 3.5 | 0.06 | - | 0.036 |
| | | | | $covs + byp + len$ | 8 | 2.8 | 0.08 | 0.006 | 0.527 |
| | | | | $covs + byp.cat + len$ | 11 | 0.1 | 0.30 | 0.010 | 0.060 |
| | | | | $covs + n.byp + len$ | 8 | 0.0 | 0.32 | 0.011 | 0.074 |
| | LGD | 71,171 | 638 | $covs$ | 6 | 20.9 | 0.00 | - | - |
| | | | | $covs + len$ | 7 | 0.1 | 0.29 | <0.001 | - |
| | | | | $covs + byp$ | 7 | 21.3 | 0.00 | - | 0.192 |
| | | | | $covs + byp.cat$ | 10 | 19.7 | 0.00 | - | 0.056 |
| | | | | $covs + n.byp$ | 7 | 19.2 | 0.00 | - | 0.053 |
| | | | | $covs + byp + len$ | 8 | 1.2 | 0.17 | <0.001 | 0.329 |
| | | | | $covs + byp.cat + len$ | 11 | 0.6 | 0.23 | <0.001 | 0.111 |
| | | | | $covs + n.byp + len$ | 8 | 0.0 | 0.31 | <0.001 | 0.147 |
| Steelhead | ULGD | 8,572 | 295 | $covs$ | 7 | 14.4 | 0.00 | - | - |
| | | | | $covs + len$ | 8 | 0.0 | 0.52 | <0.001 | - |
| | | | | $covs + byp$ | 8 | 16.1 | 0.00 | - | 0.537 |
| | | | | $covs + byp.cat$ | 11 | 20.8 | 0.00 | - | 0.800 |
| | | | | $covs + n.byp$ | 8 | 16.3 | 0.00 | - | 0.738 |
| | | | | $covs + byp + len$ | 9 | 1.3 | 0.27 | <0.001 | 0.400 |
| | | | | $covs + byp.cat + len$ | 12 | 6.5 | 0.02 | <0.001 | 0.827 |
| | | | | $covs + n.byp + len$ | 9 | 2.0 | 0.19 | <0.001 | 0.980 |
| | LGD | 29,077 | 659 | $covs$ | 6 | 116.4 | 0.00 | - | - |
| | | | | $covs + len$ | 7 | 0.1 | 0.33 | <0.001 | - |
| | | | | $covs + byp$ | 7 | 112.9 | 0.00 | - | 0.017 |
| | | | | $covs + byp.cat$ | 10 | 114.9 | 0.00 | - | 0.048 |
| | | | | $covs + n.byp$ | 7 | 112.9 | 0.00 | - | 0.019 |
| | | | | $covs + byp + len$ | 8 | 0.0 | 0.34 | <0.001 | 0.144 |
| | | | | $covs + byp.cat + len$ | 11 | 2.2 | 0.11 | <0.001 | 0.207 |
| | | | | $covs + n.byp + len$ | 8 | 0.9 | 0.22 | <0.001 | 0.271 |

**Table 3.** Simulation results: percentage of simulations with indicated outcome by species and tagging location. Tagging locations are at Lower Granite Dam (LGD) or upstream of LGD (ULGD). Models are M0 = covariates (*covs*), M1 = *covs* + number of bypass events (*n.byp*), M2 = *covs* + *n.byp* + *length*, and M3 = *covs* + *length*. Data models are those used to generate the simulated data and fitted models are those fit to the simulated data. Outcomes include a negative parameter estimate for effect of number of bypass events (*n.byp* (-)) and whether the associated *P*-value was also less than 0.05, or whether AIC for the specified fitted model was less than that from the corresponding model without *n.byp* fit to the same data. Shown for comparison are expected percentage of times each outcome would be true given the data generating model and no association between length and *n.byp*.

| Data Model | Fitted Model | Outcome | Expected | Chinook |  | Steelhead |  |
| --- | --- | --- | --- | --- | --- | --- | --- |
|  |  |  |  | ULGD | LGD | ULGD | LGD |
| M0 | M1 | *n.byp* (-) | 50.0 | 60.9 | 57.2 | 63.5 | 58.6 |
| | | *n.byp* (-) and $P < 0.05$ | 2.5 | 4.9 | 3.5 | 5.8 | 3.1 |
|  |  | *n.byp* (-) and lower AIC vs. M0 | 7.8 | 13.0 | 9.8 | 14.3 | 12.0 |
| M3 | M1 | *n.byp* (-) | 50.0 | 74.8 | 76.3 | 74.4 | 93.8 |
| | | *n.byp* (-) and $P < 0.05$ | 2.5 | 10.1 | 10.1 | 9.1 | 34.8 |
|  |  | *n.byp* (-) and lower AIC vs. M0 | 7.8 | 24.1 | 24.8 | 22.3 | 55.3 |
| M3 | M2 | *n.byp* (-) | 50.0 | 59.5 | 75.4 | 59.8 | 60.0 |
| | | *n.byp* (-) and $P < 0.05$ | 2.5 | 3.7 | 9.9 | 6.0 | 5.5 |
|  |  | *n.byp* (-) and lower AIC vs. M3 | 7.8 | 11.8 | 24.6 | 12.7 | 14.8 |

**Figure 3.**
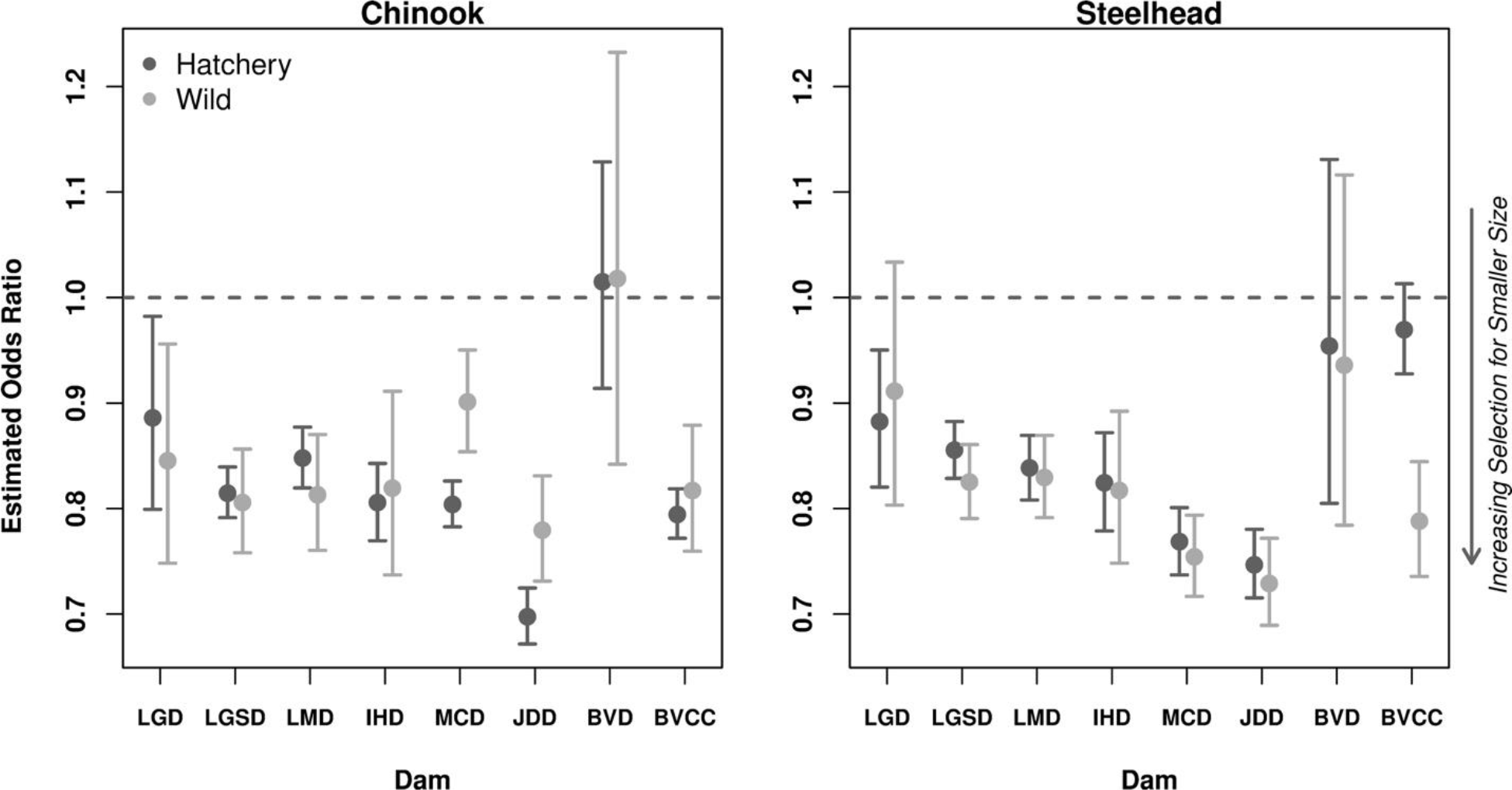
Parameter estimates and associated 95% confidence intervals for multiplicative effect of a one unit increase in standardized length on the odds of entering the bypass system at a dam. Results are shown by dam, rearing type (hatchery or wild) and species. The horizontal line at 1.0 is used as a reference to assess difference of the estimates from one. An odds ratio less than one indicates smaller fish are more likely to pass through a bypass system than pass through other routes. Abbreviations for dam names are Lower Granite Dam (LGD), Little Goose Dam (LGSD), Lower Monumental Dam (LMD), Ice Harbor Dam (IHD), McNary Dam (MCD), John Day Dam (JDD), and Bonneville Dam (BVD). BVCC is for BVD where bypass is compared to corner collector only.

**Figure 4.**
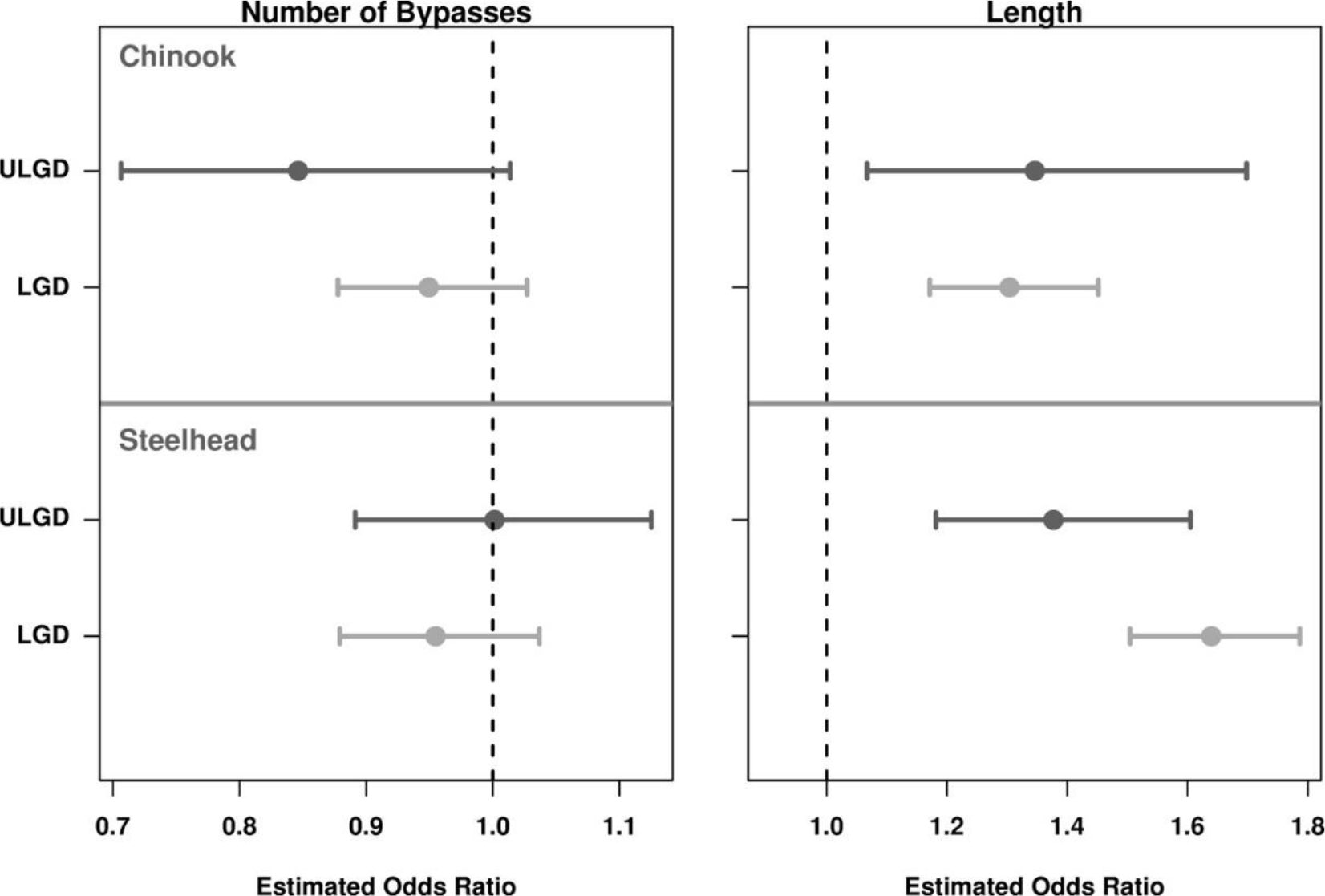
Parameter estimates and associated 95% confidence intervals for the multiplicative effects of number of bypass events and standardized length on the odds of adult return. Estimates are from models that included both number of bypass events and length. Results are shown by species and tagging location (at Lower Granite Dam (LGD) or upstream of LGD (ULGD). The vertical lines at one are used as a reference to assess difference of the estimates from one. A parameter estimate less than one for number of bypasses indicates fish with more bypass events are less likely to return as adults, and a parameter estimate greater than one for length indicates larger fish are more likely to return.

For Chinook tagged at LGD, the best model for just the covariates included fixed effects for rear type and day, and random year effects for the intercept and the slope of day. Adding length to this model reduced AIC by 20.8 with *P* < 0.0001, indicating strong evidence of an association between length and SAR. In contrast, when length was not in the model, adding the binary bypass variable increased AIC, while adding the categorical bypass variable decreased AIC by 1.2 with associated *P* = 0.056 and adding the continuous number of bypasses decreased AIC by 1.7 with *P* = 0.053. These results indicate weak to moderate evidence of an association between bypass history and SAR when not accounting for length. After length was included in the model, adding either the binary (*P* = 0.329) or categorical variables (*P* = 0.111) for bypass did not improve AIC, but adding the continuous variable for number of bypasses decreased AIC by 0.1 with *P* = 0.147. These results suggest that any association between a bypass variable and SAR could potentially be explained by length, but there is still weak evidence for a bypass effect beyond that due to length. For the model with length and continuous number of bypasses, the odds of adult return increased by an estimated multiplicative factor of 1.30 for every increase of 1 standard deviation in length (15.3 mm; 95% CI: 1.17, 1.45), and the odds of return decreased by a multiplicative factor of 0.94 for each additional bypass event (95% CI: 0.87, 1.02; Figure 3).

For steelhead tagged upstream of LGD, the best model built from only the covariates included fixed effects for rear type, tagging site, site of last detection, day, and day squared and a random year effect for the intercept. Adding length to the model resulted in a decrease in AIC of 14.4 and *P* < 0.0001, which provides strong evidence that SAR was associated with length. Adding any of the bypass variables resulted in increases in AIC whether or not length was in the model (*P* > 0.40 for each bypass variable). This suggests that there was no evidence that SAR for steelhead was associated with bypass history. For the model with length and continuous number of bypasses, the odds of adult return increased by an estimated multiplicative factor of 1.38 for every increase of 1 standard deviation in length (27.6 mm; 95% CI: 1.18, 1.61), and the odds of return increased by a multiplicative factor of 1.00 for each additional bypass event (95% CI: 0.89, 1.13; Figure 3).

For steelhead tagged at LGD, the best covariate model included fixed effects for rear type and day of year, and random year effects for the intercept and slope on day of year. Adding length to this model resulted in a reduction in AIC of 116.5 and *P* < 0.0001, which provides very strong evidence for an association between length and SAR. When length was not in the model, adding the binary bypass variable decreased AIC by 3.6 with associated *P* = 0.017, adding the categorical bypass variable decreased AIC by 1.6 with *P* = 0.048 and adding the continuous number of bypasses decreased AIC by 3.5 with *P* = 0.019. These results suggest moderate to strong evidence for an association between bypass history and SAR when not accounting for length. However, when length was in the model, adding the binary bypass variable decreased AIC by 0.1 with *P* = 0.144, while adding the categorical bypass variable increased AIC by 2.1 with *P* = 0.207 and adding the continuous number of bypasses increased AIC by 0.8 with *P* = 0.271. This indicates that after accounting for length, there was weak to no evidence for an association between bypass history and SAR remaining. For the model with length and continuous number of bypasses, the odds of adult return increased by an estimated multiplicative factor of 1.64 for every increase of 1 standard deviation in length (30.7 mm; 95% CI: 1.50, 1.79), and the odds of return decreased by a multiplicative factor of 0.95 for each additional bypass event (95% CI: 0.88, 1.04; Figure 3).

We note that rearing type was important for both species in all of the models that included length, where wild fish had higher probability of return than hatchery fish. Since hatchery fish are longer on average than wild fish, the effect of rearing type was masked and seemingly unimportant in some models without length. However, retaining rearing type in the models with length helped to more accurately estimate the length effect.

### Simulations

Our simulations show that a spurious bypass effect could arise more frequently than by chance, whether length is accounted for in the fitted model or not (Table 3). When the covariate-only model (M0) was the data-generating model, there were more negative estimates and more significant negative estimates than expected, with larger differences from expected occurring for the groups tagged upstream of LGD. This effect was likely induced by a combination of small sample sizes for groups with multiple bypass events and low overall return probabilities.

When the model with covariates and length (M3) was the data-generating model, spurious bypass effects occurred more frequently than by chance, especially when length was not accounted for in the fitted model. For the fitted model not including length (M1), a negative parameter estimate for the effect of number of bypasses occurred in greater than 74% of the simulations for each species and tagging location when no bypass effect actually existed, and a significant and negative parameter estimate occurred in at least 9% of simulations for each species and tagging location. Additionally, M1 had a lower AIC than M0 when there was also a negative estimate for bypass effect in greater than 21% of simulations for each species and tagging location. After accounting for length in the fitted model (M2), the proportion of negative estimates and the proportion of negative and significant estimates dropped but were still greater than those expected by chance. The simulation results were similar for Chinook at both tagging locations and for steelhead tagged upstream of LGD. The results for steelhead tagged at LGD indicated that a significant negative estimate of bypass effect was much more likely when length was not accounted for when compared to results for Chinook and for steelhead tagged upstream of LGD.

Mean parameter estimates across simulations were similar to parameter estimates from models fit to the observed data for all species and tagging locations except for Chinook tagged upstream of LGD (Appendix Figure A1). The estimated effect of bypass from the model fit to observed data for Chinook from upstream of LGD was much more negative than the mean of parameter estimates from the simulated data. This suggests that the association between length and number of bypass events does not completely explain the apparent effect of number of bypass events on SAR seen in the observed data.

## Discussion

We investigated associations between fish length and probability of entering juvenile bypass systems at dams, and further investigated associations among fish length, bypass history, and probability of returning as an adult for fish known to have survived through a system of hydropower dams. Our main findings were 1) there was strong evidence for a negative association between fish length and probability of bypass at most dams 2) there was strong evidence for a positive association between fish length and SAR, and 3) there was a moderate to weak evidence for a negative association between bypass history and SAR which weakened further when fish length was included in the models.

We found strong evidence for a negative association between length of fish at tagging and bypass probability for both species at six of the seven study dams, with smaller fish being more likely to enter a bypass system. There was a negative association with length at LGD for both species and rearing types, but it was not significant for wild steelhead. At BVD, there was weak to no evidence for an association with length when the bypass system was compared it to all other routes combined, but the bypass system was more likely to pass smaller fish in comparison to the corner collector alone (see discussion below). One caveat is that these two dams had much smaller sample sizes, so statistical power to detect length relationships was diminished for those data sets. The general layout of BVD is very different from the other dams in the FCRPS. The other 6 study dams are composed of a continuous structure that spans the river with a powerhouse at one end and a series of spillbays at the other. Bonneville Dam is composed of two powerhouses and a spillway which are separated by natural islands, so each is essentially in its own channel, with no direct route between powerhouse or spillway. Bonneville Dam also has the corner collector (BCC), which collects water and fish from the surface and diverts them away from the second powerhouse. These differences in dam structure could be expected to produce different fish passage behaviors that could depend on size. Finally, it should also be noted that experimental structures designed to guide fish away from the powerhouses were in periodic use at both LGD (2000, 2002, and 2006) and BVD (2008-2009), which could have affected bypass size selectivity at those sites in those years.

Our results confirm those of Zabel et al. (2005) and Hostetter et al. (2015), who found that length was an important predictor of bypass probability for both Snake River Chinook and steelhead at Snake River dams, with smaller fish more likely to enter a bypass system. Brown et al. (2013) also found that bypass probability of yearling Chinook released from LGD decreased with increasing fork length on average across multiple dams. Buchanan et al. (2011) investigated associations between bypass passage and fish length, but found mixed results with significant size selectivity evident in some release groups but not others. Similar to our study, Buchanan et al. (2011) found no relationship between size and bypass probability at LGD or BVD. Buchanan et al. (2011) did not restrict release sites to those closest to LGD, however, and did not account for time since tagging in their analyses. This likely resulted in a large number of measured lengths not representative of true lengths once fish entered the hydropower system.

Two general mechanisms that could explain size selection by bypass systems are the vertical distribution of fish as they approach a dam, and their physical ability to escape the bypass screens. The horizontal distribution of fish as they approach a dam will also affect their route of passage depending on whether they approach on the spillway side or powerhouse side, but to the best of our knowledge, this distribution is likely not dependent on fish size. The depth at which fish are swimming as they approach a dam will affect their route of passage (Li et al. 2015, 2018). Surface collection structures, such as spillway weirs, sluiceways, or the BCC, collect fish from the first few meters of the surface of the river. Entrances to standard spillbays are 10 to 15 m below the surface, while the upper ceilings of entrances to turbine intake bays are 15 to 25 m below the surface for dams on the Snake and Columbia Rivers. Bypass diversion screens extend approximately 6 to 12 m below the declining turbine intake ceilings and are designed to collect the fish orienting along the ceiling as they enter the intake. This means that fish must reach depths of approximately 20 to 35 m to escape the screens and enter the turbines. Li et al. (2015) found that yearling Chinook and steelhead that passed through juvenile bypass systems approached dams significantly deeper than those that passed though spillways, and fish that passed through turbines approached deeper than those that passed through bypass systems. If swim depth is related to size, then this could explain size differences by passage route.

Li et al. (2015, 2018) did not investigate relationships between length and swim depth but did find that subyearling Chinook, which are smaller than yearlings, traveled deeper than yearling Chinook, although they note that this could have been due to higher water temperatures when subyearling Chinook migrated in late spring and summer. It has been documented that size and level of smoltification affect buoyancy (Saunders 1965; Pinder and Eales 1969), with larger and more smolted fish being more buoyant and more likely to migrate higher in the water column. This suggests that the larger, more smolted fish are more likely to pass through spill and surface routes than through bypass or turbines. This is consistent with our findings that smaller fish were more likely to enter bypass systems compared to other routes, especially when one considers that the probability of entering turbines is low at most dams. Our results for JBS passage in comparison to BCC at BVD further support this idea, where larger fish were more likely to pass through the BCC, which is a surface route. The reason we did not find an effect of length on JBS passage at BVD when comparing to all other routes could be due to higher turbine passage at BVD. The probability of turbine passage is approximately 20% to 30% for both species at BVD (Ploskey et al. 2012), which is much higher than the other dams and results from having two powerhouses and relatively low probability of being guided by the bypass screens. If fish passing through turbines at BVD are generally smaller and those passing through spill and BCC are larger, then the combination of these groups would have a wide range of lengths, which could explain the results for JBS passage versus the other routes combined.

The second possible mechanism of size selection by bypass systems is the ability of a fish to escape when it senses the change in water velocity created by the bypass screens and gatewells (Zabel et al. 2005; Enders et al. 2012). Larger and more physically fit fish have greater strength and swim speed, which allows them a better chance to escape the bypass screens. This suggests that among fish that enter the powerhouse, fish that are guided into the bypass system would be smaller and in poorer condition on average than those that pass via turbines. However, it provides no information regarding differences in size or condition between fish that enter the powerhouse and those that pass via spill.

It was not possible in our study to distinguish whether the association between length and bypass probability was due to differential passage between the powerhouse and the spillway, or if it was driven by selection between bypass and turbine passage given entry to the powerhouse. This distinction can only be made if the exact route of passage is known for each fish, and we only had information on whether a fish passed through a bypass system or not. Further research using data from dam passage studies using radio telemetry or acoustic tags, which allow accurate determination of each route of passage, should focus on associations between length, spatial distribution, and route of passage.

Our second major finding was that there was strong evidence for an association between fish length and SAR for both species, with larger fish having higher probability of return. Many previous studies have also found length to be a significant predictor of adult return. In studies of hatchery fish, Tipping (2011) found that size at tagging had a significant effect on adult return rates in Chinook in 7 of 10 release groups. Releases of hatchery fish in the Deschutes River (a tributary that enters the Columbia River upstream of two FCRPS dams) also showed length to be a strong predictor of adult return (Beckman et. al 1999). Cross et al. (2009) found a significant effect of length on the marine survival of pink salmon. Evans et al. (2014) found juvenile length and condition to be strong predictors of adult survival in steelhead from the Columbia and Snake Rivers. A mass review of marine survival studies of anadromous salmonids spanning a range of species and tag types found fish length to be one of the most frequent significant predictors of fish survival (Drenner et al., 2012). In a study of Chinook from the Willamette River (a Columbia River tributary that enters downstream of the FCRPS) based on scale analysis, Clairborne et al. (2011) found a significant effect of ocean entry size on adult survival in 3 of four years examined. However, Romer et al. (2013) did not find a significant effect of fish length on adult survival and return in two costal groups of steelhead tagged in 2009. Zabel and Williams (2002) found larger yearling Chinook were more likely to return as adults, and Zabel et al. (2005) and Passolt and Anderson (2013) reported a size-dependent survival pattern in the FCRPS.

Three mechanisms controlling size-selective mortality in teleost fish were suggested by Sogard (1997): differences in vulnerability to predation, susceptibility to starvation, and tolerance of environmental extremes. Size-specific consumption by predators can be due to limitations of mouth gape size, behavioral selection, or size-dependent escape ability of the prey. Size-selective predation on juvenile salmonids has been documented in the early phase of ocean residence (Parker 1971; Healy 1982; Hargraves and LeBrasseur 1986; Holtby et al. 1990). As predators themselves, larger salmon smolts have a wider selection of prey available due to their larger gape sizes and faster swim speeds, allowing faster growth in the ocean and lower probability of starvation. Larger fish also mature faster and return at earlier ages (Scheuerell 2005; Tattam et al. 2015), which means they have less time exposed to mortality risks in the ocean.

Our third major finding was that there was moderate to weak evidence for a negative association between bypass history and probability of adult return, but that association weakened when length was accounted for. For Chinook tagged at LGD, and steelhead tagged at or upstream of LGD, bypass variables were insignificant without length or became insignificant when length was also in the model. Only for Chinook tagged upstream of LGD was number of bypass events marginally significant after accounting for length.

The negative association between fish length and bypass probability was the most likely explanation for the cases where evidence for a bypass effect on SAR diminished or disappeared after accounting for fish length. Smaller fish were more likely to experience multiple bypasses, so the number of bypasses essentially functioned as a surrogate for length in the model. It is clear from our models of bypass passage that smaller fish have a higher probability of bypass passage at most dams when compared to larger fish, which translates into a higher expected number of bypass events during migration. If length is not explicitly accounted for in a model for adult return, inclusion of a variable for number of bypass events (or even for number of spillway passage events) in the model could act as a surrogate for fish length. If the bypass variable does not contain additional explanatory power beyond that offered by the correlation with length, then one would expect the apparent effect of the bypass variable on SAR to disappear when length is included in the same model. We saw this phenomenon occur for both steelhead and Chinook tagged at LGD. Our simulations further indicated that a false signal for number of bypasses could be detected as significant more often than by chance when there is no true association with SAR due to the association between length and probability of entering bypass systems.

When evidence for a negative effect of bypass on SAR remains after accounting for length and other confounding variables, it suggests there could be lasting effects of multiple bypass events on fish. Our results for Chinook tagged upstream of LGD support this possibility. Passage through bypass systems at multiple dams could be causing an accumulation of trauma and stress that results in impaired condition, reduced energy reserves, and increased susceptibility to predation in the estuary and ocean, as has been suggested by others (e.g., Budy et al. 2002; Schaller et al. 2013). Although this seems like a reasonable assumption biologically, the available direct empirical evidence in support of this is mixed.

Maule et al. (1988) found blood measures of stress increased cumulatively as fish passed through points in the JBS at McNary Dam, and several studies summarized by Ferguson et al. (2005) showed increased indices of stress in bypass systems at other dams, but indices returned to pre-stress levels in a relatively short time. Barton et al. (1986) found multiple handling events of juvenile Chinook resulted in cumulative physiological stress. Barton and Schreck (1987) found a relationship between multiple stress events and increased metabolic rate in juvenile steelhead, suggesting that multiple stress events, such as multiple bypasses, could result in decreased energy reserves. Mesa (1994) exposed juvenile Chinook to multiple stressors and found preferential predation by northern pikeminnow (*Ptycholcheilus oregonensis*), an important predator of salmonids in the FCRPS (Rieman et al. 1991), on the stressed individuals compared to controls in a short period, but no differences in predation after one hour of recovery. Sandford et al. (2014) investigated the delayed effect of bypass passage on post-hydropower system survival by collecting juvenile Chinook at BVD and holding them in seawater tanks and found no effect of bypass history on survival, but this study could not account for predation or factors that occur in the natural environment.

The proportion of yearling Chinook and steelhead experiencing some level of descaling due to JBS passage ranged from 1.5 to 9.6% in studies of bypass passage at individual dams in the FCRPS (Ferguson et al. 2005). Multiple bypass events would certainly increase the probability that a fish experiences some level of descaling or trauma. Evans et al. (2014) found that steelhead with general bodily injuries had higher susceptibility to avian predation but level of descaling was not influential. Gadomski et al. (1994) found that experimentally descaled juvenile Chinook had short-term physiological stress responses but were not more susceptible to predation than controls. Descaled juvenile salmon can suffer high mortality due to impaired ability to osmoregulate when exposed to seawater within 1 day of the descaling event, but can recover and survive at normal levels if allowed to remain in fresh water for a few days (Bouk and Smith 1979, Zydlewski et al. 2010). Juveniles migrating the FCRPS would have sufficient time to recover before reaching the estuary.

It is difficult to explain why multiple bypass events would affect SAR more for Chinook tagged upstream than those tagged at LGD, and why they would affect Chinook but not steelhead, especially when we consider that Chinook from upstream of LGD had the smallest sample size and least number of returning adults compared to the other groups. We cannot rule out the possibility that the results for Chinook from upstream of LGD were a spurious relationship driven by small sample sizes. The direction of the bypass effect for Chinook tagged upstream of LGD was not consistent from year to year, based on the direct SAR estimates for each number of bypass events (see Appendix). We were also not able to account for all confounding variables that are associated with both bypass probability and adult return, such as fish condition and disease status, which may have differed among the tagging locations. Given these uncertainties, the evidence for a causal relationship between number of bypass events and SAR for Chinook tagged upstream of LGD is still questionable.

Another important point is that the probability of experiencing a particular number of bypass events decreases rapidly for each additional number of events greater than two (Appendix Table A2). This means that the proportion of the migrating populations of yearling Chinook and steelhead experiencing more than three bypass events is low. Our results indicated that SAR for fish with zero to two or three bypass events tended to be similar (Appendix Figures A2-A4). Therefore, the overall SARs for these migrating populations would not be affected much by any delayed mortality experienced by multiply bypassed fish.

We were only able to investigate effects of passing through bypass systems and were not able to investigate effects of passing through turbines or other routes due to lack of PIT tag detection in those routes. Turbine passage can result in rapid pressure changes and strikes with blades and other structures in the turbine housing. Multiple passage events through turbines would be expected to lead to accumulated trauma and stress and increased susceptibility to predation. Ferguson et al. (2007) found that juvenile salmon had significant delayed mortality between 15 and 46 km downstream after passing through turbines at McNary Dam and concluded the likely cause was impaired sensory systems that led to increased vulnerability to predation. Studies on the delayed effects of turbine or other routes of passage on survival beyond the hydropower system are needed but are lacking, mostly due to limitations of tagging technologies.

Others have attempted to estimate effect of passage through either JBS or turbines on survival to adulthood by creating an index of powerhouse passage for release groups of fish (Petrosky and Schaller 2010; Schaller et al. 2013; McCann et al. 2017). There are different ways of calculating these indices based on various assumptions about passage route probabilities, but the resulting group-level metric is an estimate of the expected number of powerhouse passage events experienced by each fish across the set of dams passed during the migration. These methods do not account for the known bypass events of individual fish and there is no way to know which fish actually went through turbines. There is also no way to know if the individual fish that actually went through turbines or JBS had lower probability of return. These authors found negative associations between survival to adulthood and the indices of number of powerhouse passages, but they did not account for fish size or condition in their models. Even if they would have included fish characteristics, modeling approaches like this that use temporal or spatial aggregations of fish as observational units can only account for individual fish characteristics as group-level summary statistics, which results in loss of important information. There is a need to be able to account for individual fish characteristics as well as individual passage route histories in our modeling of survival to adulthood, but doing so directly is not possible given current data limitations.

Ideally, we would have tag detection in every route of passage and could then explicitly link actual route of passage to the fate of individual fish. Monitoring of PIT tags in this way is costly and not a viable option in the near future. Active (radio or acoustic) tags allow determination of route of passage with fairly high precision, but the current life of those tags is short and cannot be used to obtain data on adult returns. Fish tagged with both PIT and active tags could provide solutions, but such tagging efforts are costly and burdensome on the fish, resulting in small sample sizes and results potentially not representative of the population at large. A secondary solution could be to develop more sophisticated models that would take advantage of all available tagging data and would account for the unknown passage routes (spill or turbine) of individual fish using probabilistic relationships that depend on fish length and other covariates, such as in a Bayesian framework. Such models may offer more accurate predictions of effects of specific routes of passage on adult returns than are currently available. This is an area of our current research.

In conclusion, based on our results and those of others, it is imperative that researchers investigating SAR or bypass probability include length of individual fish in their models. If other data related to measures of individual fish health exist, such as condition factor, disease status, etc., then those data should also be included. Neglecting these important sources of information could lead to spurious modeling results which could misinform management decisions and lead to misallocation of limited resources.

## Acknowledgements

Mark Scheuerell and Steve Smith provided valuable reviews that improved the manuscript. Marvin Shutters provided information about deployment of the behavioral guidance structure at Lower Granite Dam. We thank all of the agencies, organizations, and individuals involved in collecting and tagging the PIT-tagged fish we used in this study, and we thank PTAGIS for housing, managing, and making publically available all of the PIT tag data.

## Appendix

### Data Summaries

Here we provide some summaries of the data used in the SAR models. Table A1 shows the relationship between lengths at tagging, some measures of bypass passage, and percentage of adults returning, with larger fish having fewer bypass events and higher SAR. Table A2 provides information on length and on percentage of fish experiencing different number of bypass events. The majority of fish experienced between zero and two bypass events, and the proportions decrease rapidly for each successive increase in number of bypasses. Only 3-15% experienced four or more bypass events, with results depending on species, rearing type, and tagging location.

**Table A1.**
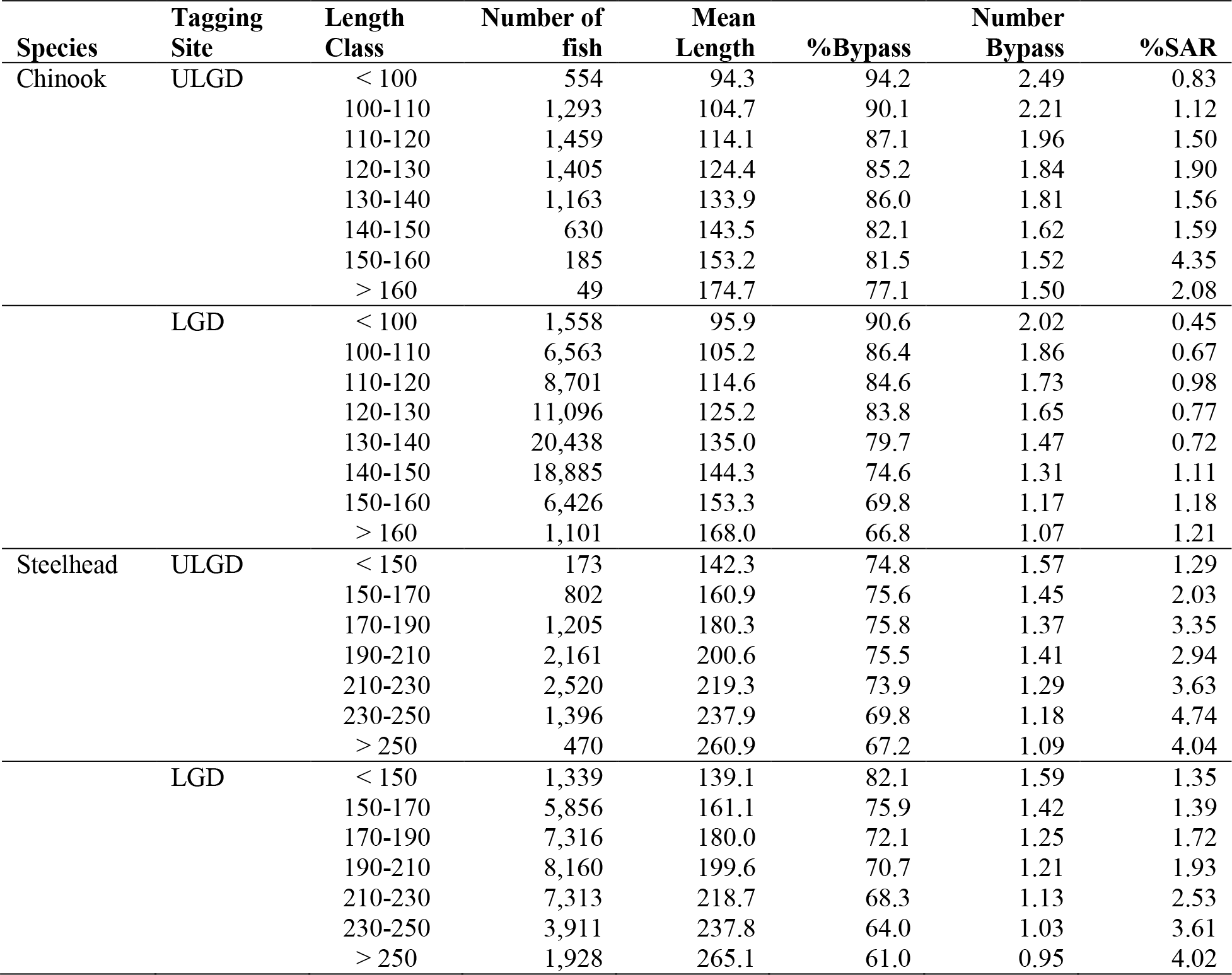
Summary of observed data used in SAR models (2004-2014), by species, tagging location, and length class (mm). Shown are number of fish detected as juveniles, mean length (mm), percentage of fish with at least one bypass event (%Bypass), average number of bypass events (Number Bypass), and percentage returning as adults (%SAR). SAR here is calculated as the number of returning adults divided by the number of juveniles within each length class. Tagging locations are Lower Granite Dam (LGD) and upstream of LGD (ULGD). Fish tagged at LGD have one fewer potential bypass location compared to those tagged ULGD.

**Table A2.** Summary of data on length at tagging (mm) and percentage of fish with each possible number of bypass events by species, rear type (hatchery (H) or wild (W), and tagging site (at Lower Granite Dam (LGD) or upstream of LGD) for 2004-2014. Number of bypass events for fish tagged at LGD are those experienced at dams downstream of LGD, so maximum number of bypasses is 7 for ULGD fish and 6 for LGD fish.

| Species | Site | Type | Length |  |  | Number of Bypass Events |  |  |  |  |  |  |
| --- | --- | --- | --- | --- | --- | --- | --- | --- | --- | --- | --- | --- |
|  |  |  | N | Mean | SD | 0 | 1 | 2 | 3 | 4 | 5 | ≥6 |
| Chinook | ULGD | H | 3,071 | 132 | 13 | 14 | 30 | 28 | 18 | 8 | 2.0 | 0.3 |
|  |  | W | 3,277 | 111 | 11 | 13 | 26 | 27 | 20 | 11 | 3.3 | 0.8 |
|  | LGD | H | 58,059 | 138 | 11 | 22 | 35 | 26 | 12 | 4 | 0.6 | 0.1 |
|  |  | W | 13,112 | 110 | 8 | 15 | 30 | 29 | 18 | 7 | 1.5 | 0.1 |
| Steelhead | ULGD | H | 6,490 | 218 | 21 | 26 | 36 | 23 | 10 | 4 | 0.9 | 0.1 |
|  |  | W | 2,082 | 177 | 22 | 26 | 35 | 22 | 11 | 4 | 1.2 | 0.5 |
|  | LGD | H | 18,338 | 216 | 26 | 31 | 37 | 21 | 8 | 2 | 0.4 | 0.0 |
|  |  | W | 10,739 | 178 | 23 | 28 | 35 | 23 | 10 | 3 | 0.7 | 0.1 |

### Detailed Model Selection Results

Here we present some additional information about the models selected for the bypass probability modeling and the SAR modeling. Table A3 shows the covariate terms in the best models for bypass probability before inclusion of length at tagging, and Table A4 shows the forms of the top three covariate-only models for probability of adult return ranked by AIC.

**Table A3.**
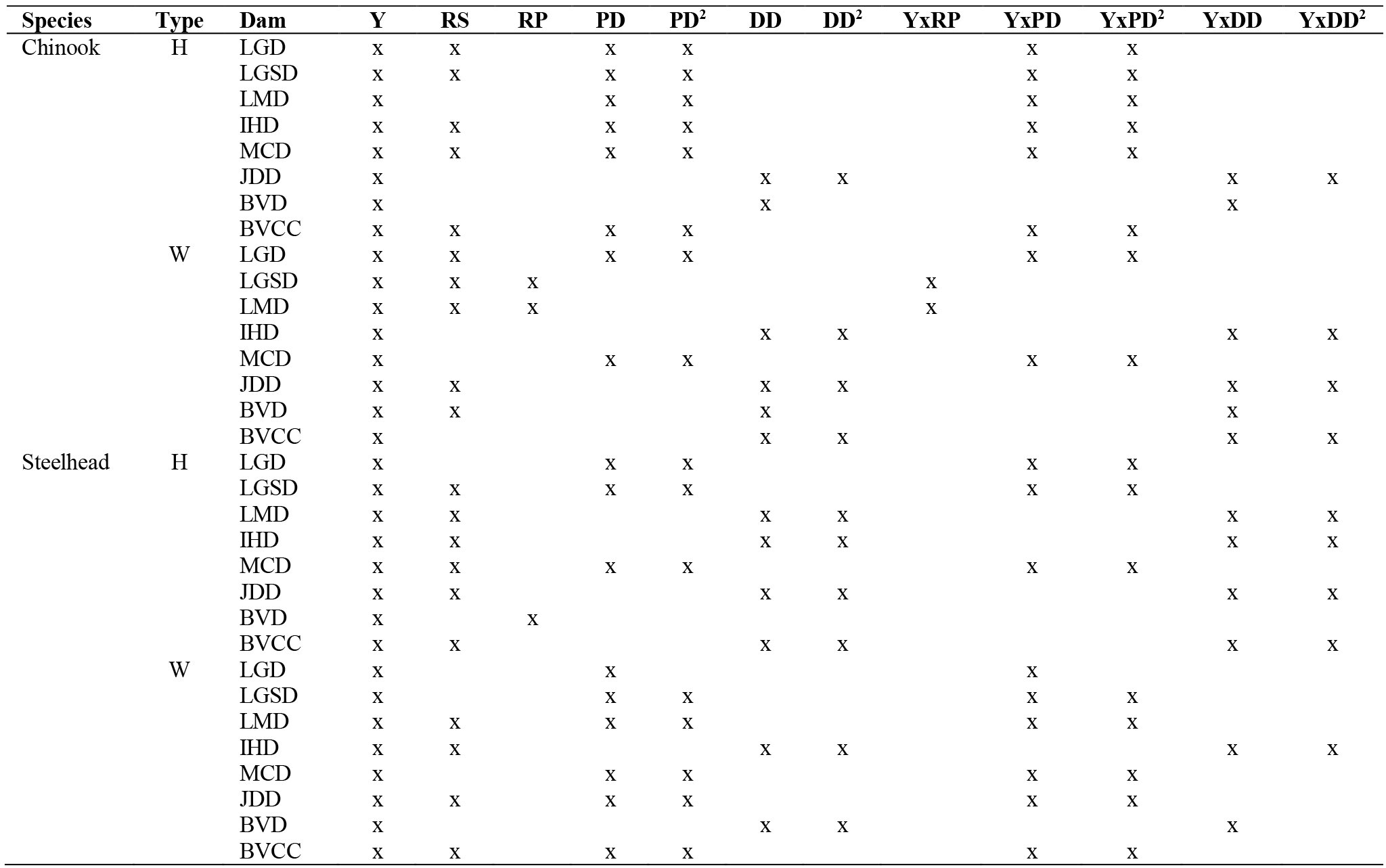
Best models for bypass probability before inclusion of length at tagging (models with covariates only). Models are shown by species, rear type (Type) and dam. Possible main variables are year (Y), release site (RS), release period (RP), passage day at LGD (PD), squared passage day (PD^2^), detection day (DD) at Bonneville Dam or the Estuary Towed Array, and squared detection day (DD^2^). Interactions between year and other variables are preceded by Yx. Variables included in the best models are indicated by an x.

**Table A4.**
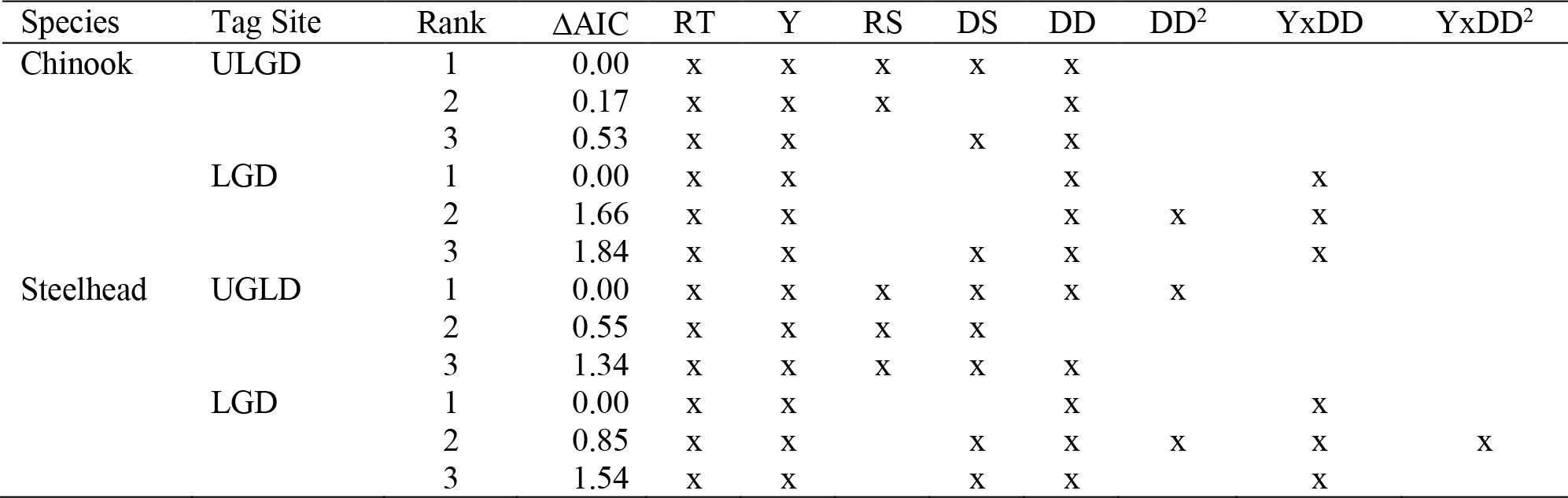
Top three models for probability of adult return before inclusion of length at tagging or bypass variables (models with covariates only) by species and tagging site. Tagging sites are at Lower Granite Dam (LGD) or upstream of LGD (ULGD). Model rank and difference in AIC (ΔAIC) compared to the model with lowest AIC are shown for each model. Possible model variables are rear type (RT), year (Y), release site (RS), site of late detection (DS), detection day (DD), squared detection day (DD^2^), and interactions between year and DD (YxDD) and year and DD^2^ (YxDD^2^). Models for LGD do not include RS as a possible variable. Inclusion of a variable in a model is indicated by an x.

### Simulation Details

We simulated data by first drawing a set of model parameter values from a multivariate normal distribution where the mean was the vector of parameter estimates (both fixed and conditional random effects) from the best model with length and covariates (but no bypass effect) fit to the real data and the covariance matrix was the estimated joint covariance of the associated model parameters. What follows is a more detailed description of the general model and covariance structure.

Suppose we have a data set with *n* observations for which we are going to apply a generalized linear mixed model with *p* fixed effects and *q* random effects. The model equation for the predicted (mean) values on the scale of the link function can be written in matrix notation as

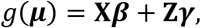

where *g*(·) is the link function, ***μ*** is an *n* × 1 vector of predicted values on the observation scale, ***X*** is an *n* × *p* design matrix for the fixed effects, ***β*** is a *p* × 1 vector of parameters for the fixed effects, ***Z*** is an *n* × *q* design matrix for the random effects, and ***γ*** is a *q* × 1 vector of random effects. The maximum likelihood estimates 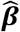 for the fixed effect parameters are assumed to follow a multivariate normal distribution with mean ***β*** and covariance matrix (**X′WX**)^−1^, where **W** is a *n* × *n* diagonal weight matrix with *W*_*ii*_ = *μ*_*i*_(1 − *μ*_*i*_), for *i* = 1, …, *n*, for the logistic regression with logit link. In general, the random effects are assumed to follow a multivariate normal distribution with an *n* × 1 vector of zeros as the mean and *q* × *q* covariance matrix **G**. The covariance of the estimated set of conditional random effects 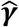 is (**Z′WZ** + **G**^−**1**^)^−1^. It follows that the joint covariance matrix of the fixed effect parameters and the conditional random effects is:

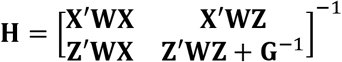

For drawing new values of the fixed and random parameters, we used a multivariate normal distribution with mean based on the 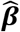 and 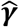 estimated from the fit to the original data and covariance matrix 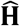, where 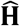 is just **H** with the elements of **W** based on the predicted values 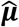, and the component **G** replaced by the estimated 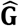 from the original model fit. See McCulloch (2003) or Bates et al. (2015) for more information on generalized linear mixed model theory.

### Additional Simulation Results

Here we use a set of plots to summarize the parameter estimates from the simulation studies and compare them to the estimates from corresponding models fit to the observed data.

**Figure A1.**
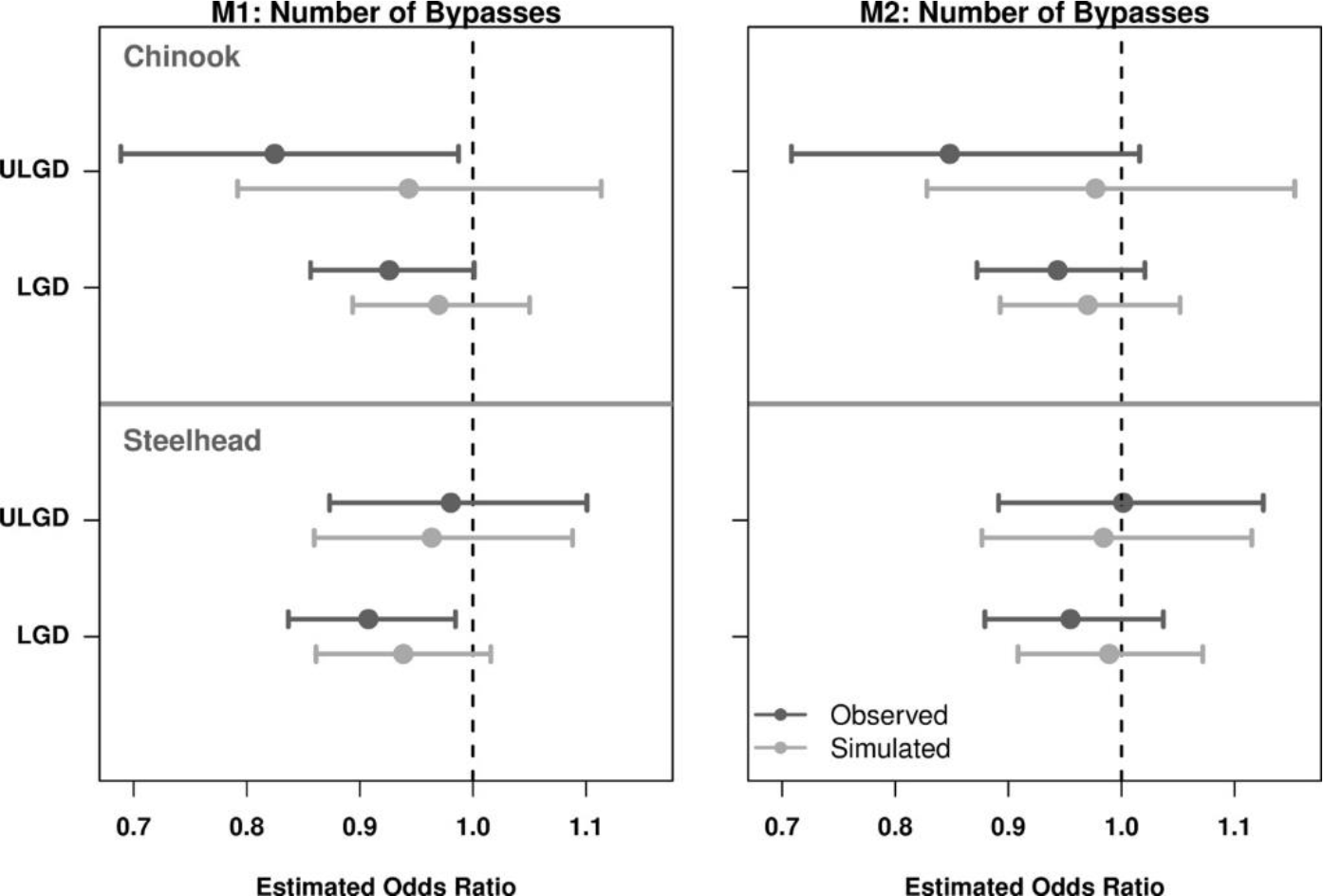
Parameter estimates for effect of number of bypass events for model M1 = covariates (*covs*) + number of bypass events (*n.byp*) and model M2 = *covs* + *n.byp* + *length* for models fit to observed data and models fit to simulated data. The data generating model for the simulated data sets was model M3 = *covs* + *length*. Interval estimates are 95% confidence intervals for the estimates from models fit to observed data, and are 0.025 and 0.975 quantiles of estimates from models fit to simulated data. Point estimates from simulated data fits are means across simulations.

### Annual Variation in SAR and Effects of Sample Size

To investigate annual variation in the association between adult return probability and number of bypass events and to investigate effects of small sample sizes, we calculated empirical return proportions for each number of bypass events in each year for both species and tagging locations. We then fit simple SAR models to the entire data sets of individual fish (as were used in the main analyses) with fixed effects for number of bypass events and rearing type and random effects for year. Models were fit separately by species and tagging location. Predicted probabilities within each year were calculated using the model formulas and intercepts were adjusted according to the proportion of wild fish vs. hatchery fish within each year. Predicted probabilities for all years combined were calculated by weighting the random year effects by the proportion of fish in each year and weighting the rearing type coefficient by the proportion of wild fish in each year. These predicted probabilities were calculated for visual comparison to the empirical SAR values (see Figures A1-A4). An annual empirical SAR was also calculated for each year and plotted as a horizontal line, and the number of juveniles and returning adults for each year and number of bypass events are displayed on each plot.

A few patterns are evident across the different species and tagging locations. In most years there are so few juveniles with four or more bypasses (and sometimes three or more), that even if we assume constant (not decreasing) return probabilities across all bypass events, the expected number of returning adults is less than one, where the expected value is the return probability multiplied by the number of juveniles in each group. The resulting number returning is therefore often zero and the empirical SAR is also zero, which is an underestimate of the true return probability.

Another pattern to notice is that often the empirical estimates for zero to two or three bypass events are scattered around the annual mean SAR and do not show a decreasing pattern. This suggests that return probabilities are equal for zero to three bypass events. Unfortunately, there are often too few juveniles and therefore too few returns for four or more bypass events (and sometimes three or more) to get a reliable estimate of SAR. Even when all years are combined, there are often too few juveniles with five or more bypass events to expect non-zero returns.

**Figure A2.**
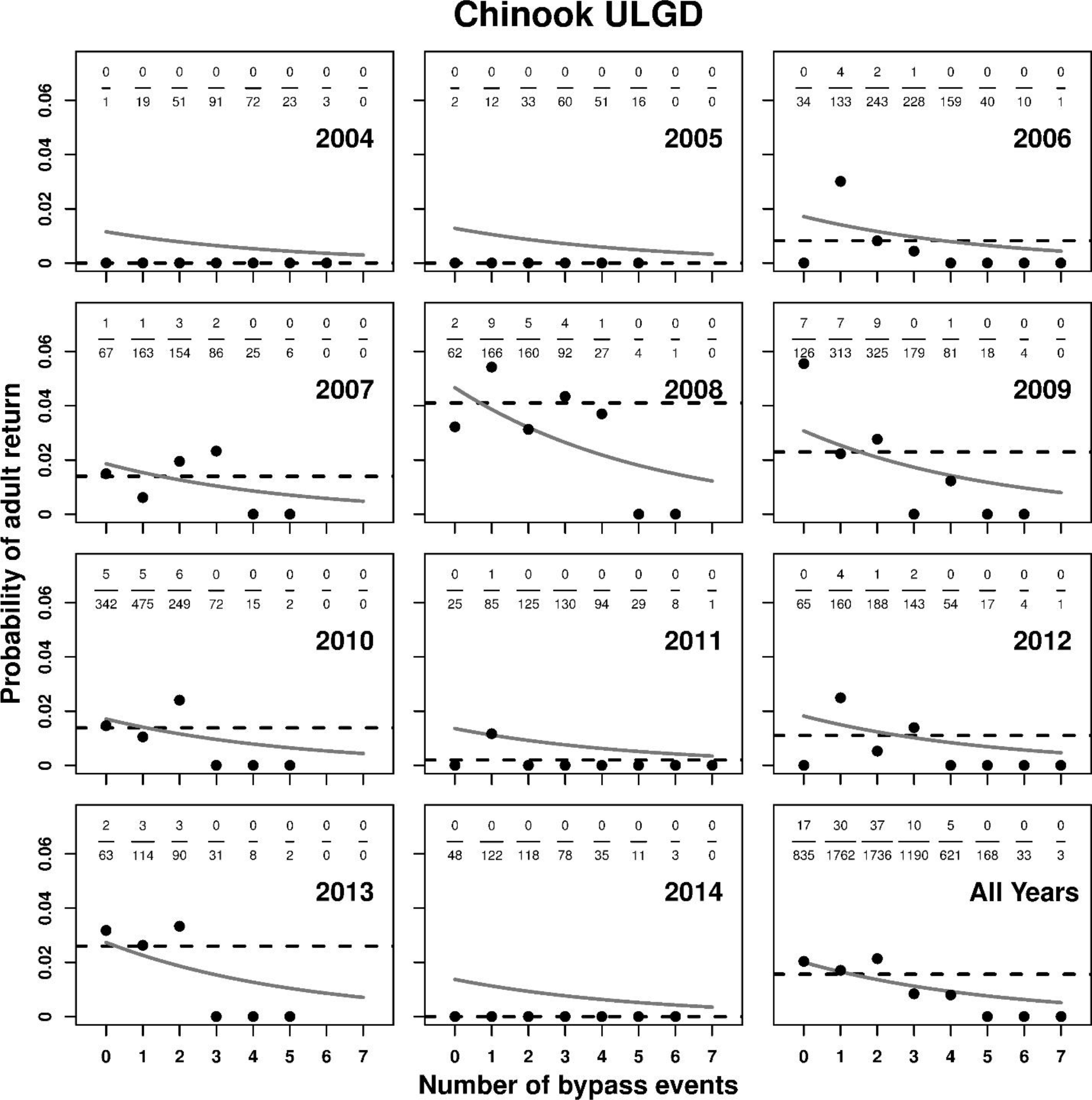
Probability of adult return (SAR) by number of bypass events and year for Chinook tagged upstream of Lower Granite Dam (ULGD). Also shown are results for all years combined. For each year and number of bypass events the number of juveniles (denominator) and number of returning adults (numerator) are shown as fractions at the top of each panel. Black circles are the corresponding empirical SAR estimates, horizontal dashed lines are annual empirical SAR estimates for all bypass categories combined, and gray lines are predicted probabilities from SAR models with fixed effects for rearing type and number of bypass events and random effects for yearly intercepts.

**Figure A3.**
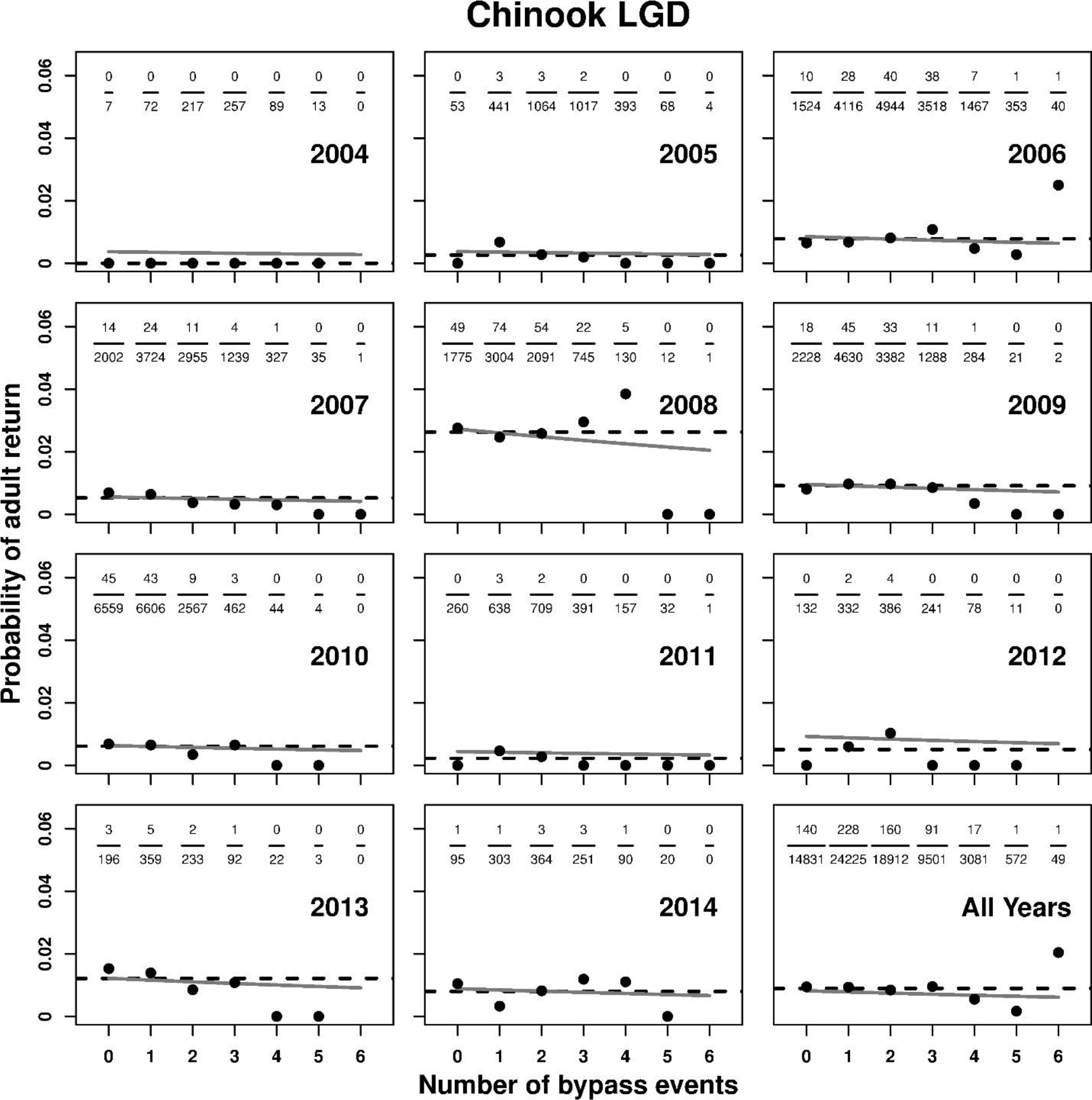
Probability of adult return (SAR) by number of bypass events and year for Chinook tagged at Lower Granite Dam (LGD). Also shown are results for all years combined. For each year and number of bypass events the number of juveniles (denominator) and number of returning adults (numerator) are shown as fractions at the top of each panel. Black circles are the corresponding empirical SAR estimates, horizontal dashed lines are annual empirical SAR estimates for all bypass categories combined, and gray lines are predicted probabilities from SAR models with fixed effects for rearing type and number of bypass events and random effects for yearly intercepts.

**Figure A4.**
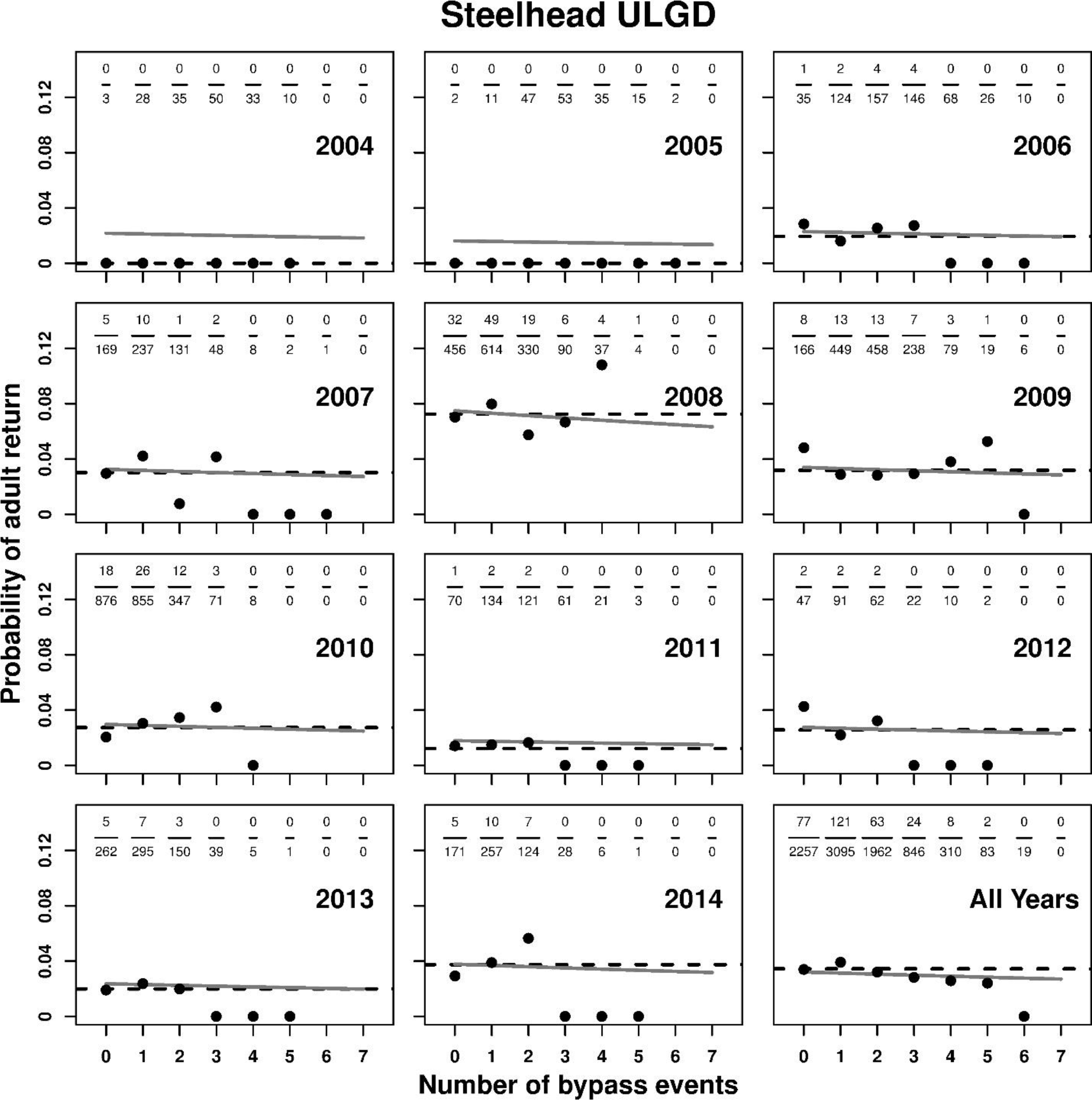
Probability of adult return (SAR) by number of bypass events and year for steelhead tagged upstream of Lower Granite Dam (ULGD). Also shown are results for all years combined. For each year and number of bypass events the number of juveniles (denominator) and number of returning adults (numerator) are shown as fractions at the top of each panel. Black circles are the corresponding empirical SAR estimates, horizontal dashed lines are annual empirical SAR estimates for all bypass categories combined, and gray lines are predicted probabilities from SAR models with fixed effects for rearing type and number of bypass events and random effects for yearly intercepts.

**Figure A5.**
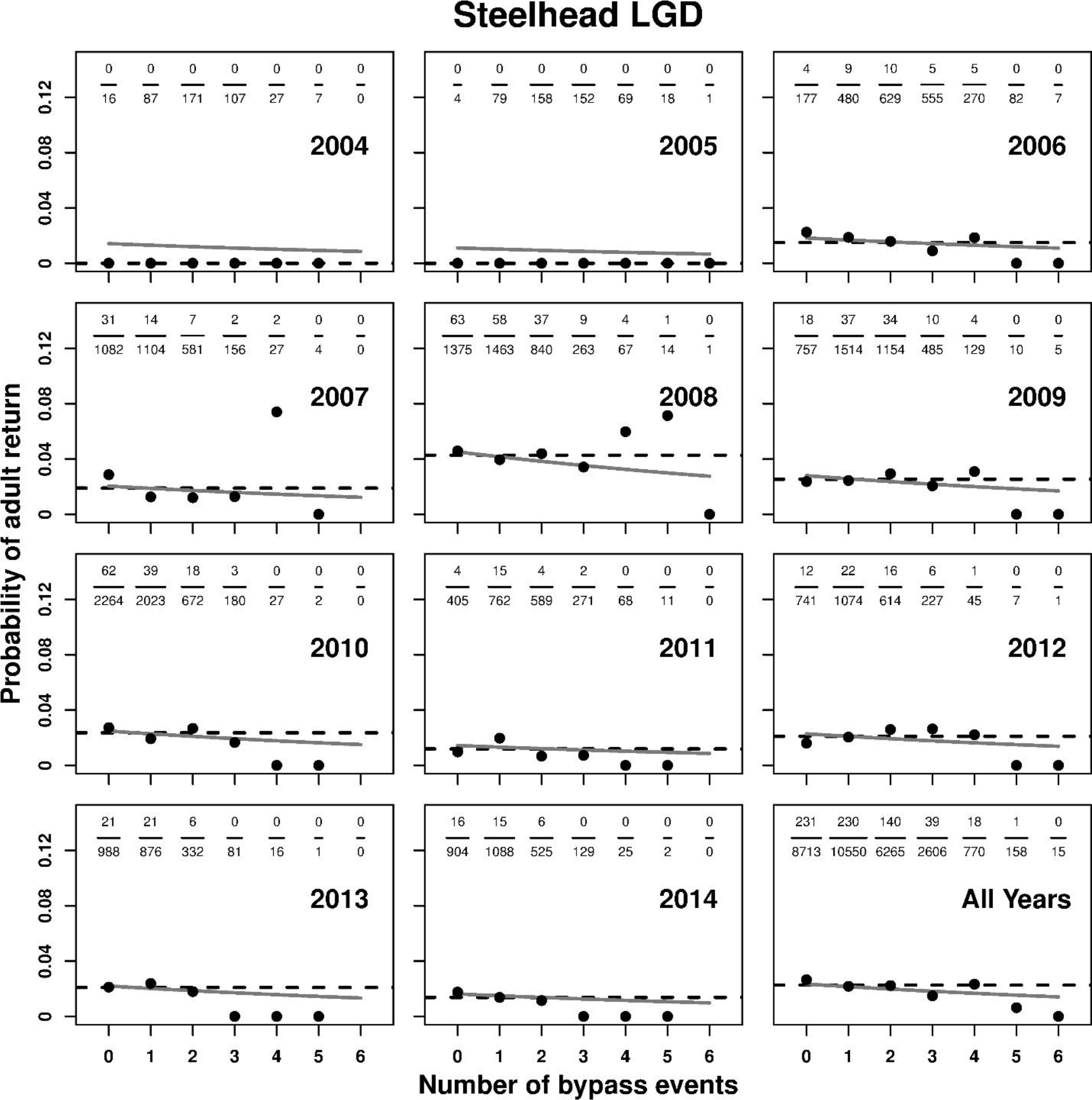
Probability of adult return (SAR) by number of bypass events and year for steelhead tagged at Lower Granite Dam (LGD). Also shown are results for all years combined. For each year and number of bypass events the number of juveniles (denominator) and number of returning adults (numerator) are shown as fractions at the top of each panel. Black circles are the corresponding empirical SAR estimates, horizontal dashed lines are annual empirical SAR estimates for all bypass categories combined, and gray lines are predicted probabilities from SAR models with fixed effects for rearing type and number of bypass events and random effects for yearly intercepts.

